# Scalable single-cell metagenomic analysis with Bascet and Zorn

**DOI:** 10.1101/2025.06.20.660799

**Authors:** Hadrien Gourlé, Iryna Yakovenko, Jyoti Verma, Julian Dicken, Florian Albrecht, Arthur Stenlund, Chetan Sharaf, Chinmay Dwibedi, Linas Mažutis, Johan Normark, Nongfei Sheng, Nicklas Strömberg, Tommy Löfstedt, Laura M. Carroll, Johan Henriksson

## Abstract

Single-cell metagenomic sequencing (scMetaG) can provide maximum-resolution insights into complex microbial communities. However, existing bioinformatic tools are not equipped to handle the massive amounts of data generated by high-throughput scMetaG methods. Here, we present a bioinformatic toolkit for end-to-end scMetaG analysis: Bascet, a command-line suite designed to scale to massive scMetaG datasets (≥1 million cells); and Zorn, an R package/workflow manager that enables reproducible scMetaG data analysis, exploration, and visualization (http://zorn.henlab.org/). Enabled by recent advances in droplet microfluidics, we use Bascet and Zorn to develop and optimize a high-throughput scMetaG method on a ten-species mock community. To showcase their utility on a clinically relevant sample, we apply Bascet and Zorn to human saliva, generating single-amplified genomes from >15k prokaryotic cells, including uncultured, low-abundance taxa undetected by bulk metagenomics. Overall, Bascet and Zorn enable reproducible scMetaG analysis, allowing users to query microbiomes at unprecedented resolution and scale.

## INTRODUCTION

Determining “who’s there” in a microbial community (e.g., which species/strains are present; which genes they carry) remains a fundamental, yet non-trivial question in microbiome research.^1,2^ To this end, single-cell metagenomic sequencing (scMetaG) methods can provide maximum-resolution insights into complex microbial communities.^1,3,4^ Also known as single-cell whole-genome sequencing (scWGS) or single-cell DNA-sequencing (scDNAseq), scMetaG generates sequencing reads, which are linked to their microbial cell of origin. Because the single-cell origin of each sequencing read is known, scMetaG can uncover heterogeneity, which bulk methods like shotgun metagenomics lack the resolution to capture (e.g., by differentiating closely related strains; by identifying pan-genomic differences among cells).^1,3^

Until recently, scMetaG has been carried out in a multiwell (e.g., 96-well) plate format,^5^ making it low-throughput and prohibitively expensive. However, droplet scMetaG methods, with the potential to query millions of cells per sample, have recently emerged.^3,6,7^ Droplet scMetaG methods encapsulate single bacterial cells in e.g., oil-in-water emulsion droplets (e.g., 10x Genomics)^8^, or more recently, semi-permeable capsules (SPCs; e.g., Atrandi Biosciences),^9,10^ aiming to make scMetaG scalable and economically feasible.

Should it be possible to scale existing droplet scMetaG methods to their theoretical limits, microbiologists would face substantial challenges analyzing the resulting data. Currently, there is no dedicated software available for scMetaG analysis, and existing bioinformatic tools are not equipped to handle this novel data modality. For example, all software for eukaryotic single-cell analysis (e.g., scRNAseq, spatial transcriptomics) rely on a known reference genome and thus cannot be used for scMetaG. Likewise, methods for bulk metaG/bacterial WGS analysis cannot scale to the massive amounts of data that droplet scMetaG methods have the potential to generate without major modifications. For example, in the scRNAseq space, commercial kits have the capacity to generate 10^6^ single-cell transcriptomes per run; for comparison, there are 10^6^ total assembled bacterial genomes in the NCBI Assembly database (https://www.ncbi.nlm.nih.gov/assembly/, accessed 25 September 2026).^11^ Thus, existing single-cell and bulk analysis software are not designed for the throughput enabled by droplet scMetaG methods.

Here, we present Bascet and Zorn, a bioinformatic toolkit for scalable, reproducible, end-to-end scMetaG analysis. We describe the overall design of Bascet and Zorn and outline major steps in a typical analysis workflow. Enabled by recent advances in droplet microfluidics, we use Bascet and Zorn to optimize a novel, high-throughput, SPC-based scMetaG method on a mock community. Finally, to showcase their utility on a clinically relevant sample, we use Bascet and Zorn to analyze scMetaG data from a human saliva sample, generating single-amplified genomes (SAGs) from >15k prokaryotic cells, including uncultured, low-abundance taxa undetected by bulk metagenomics. Overall, Bascet and Zorn enable reproducible scMetaG analysis, allowing users to query microbiomes at unprecedented resolution and scale.

## RESULTS

### Bascet and Zorn enable scalable processing of single-cell data

Recent single-cell and droplet microfluidics advances indicate that it is now realistic to generate scMetaG libraries for up to 10^6^ prokaryotic cells in a single run (**Fig 1a**). However, existing bioinformatic tools for bacterial WGS analysis are not designed to scale to massive amounts of this novel type of data. For example, cluster job managers do not cope well with many small jobs, and small files subject modern file systems to excessive performance penalties (**Supplemental Note**). Naively partitioning paired-end reads from each cell in one scMetaG sample with 10^6^ cells would produce 2·10^6^ files for paired-end read storage alone; if one desires to perform additional analyses (e.g., SAG assembly, quality control [QC], annotation), the number of files required to store this data increases rapidly, hindering the performance of most modern filesystems and software.

**Figure 1.**
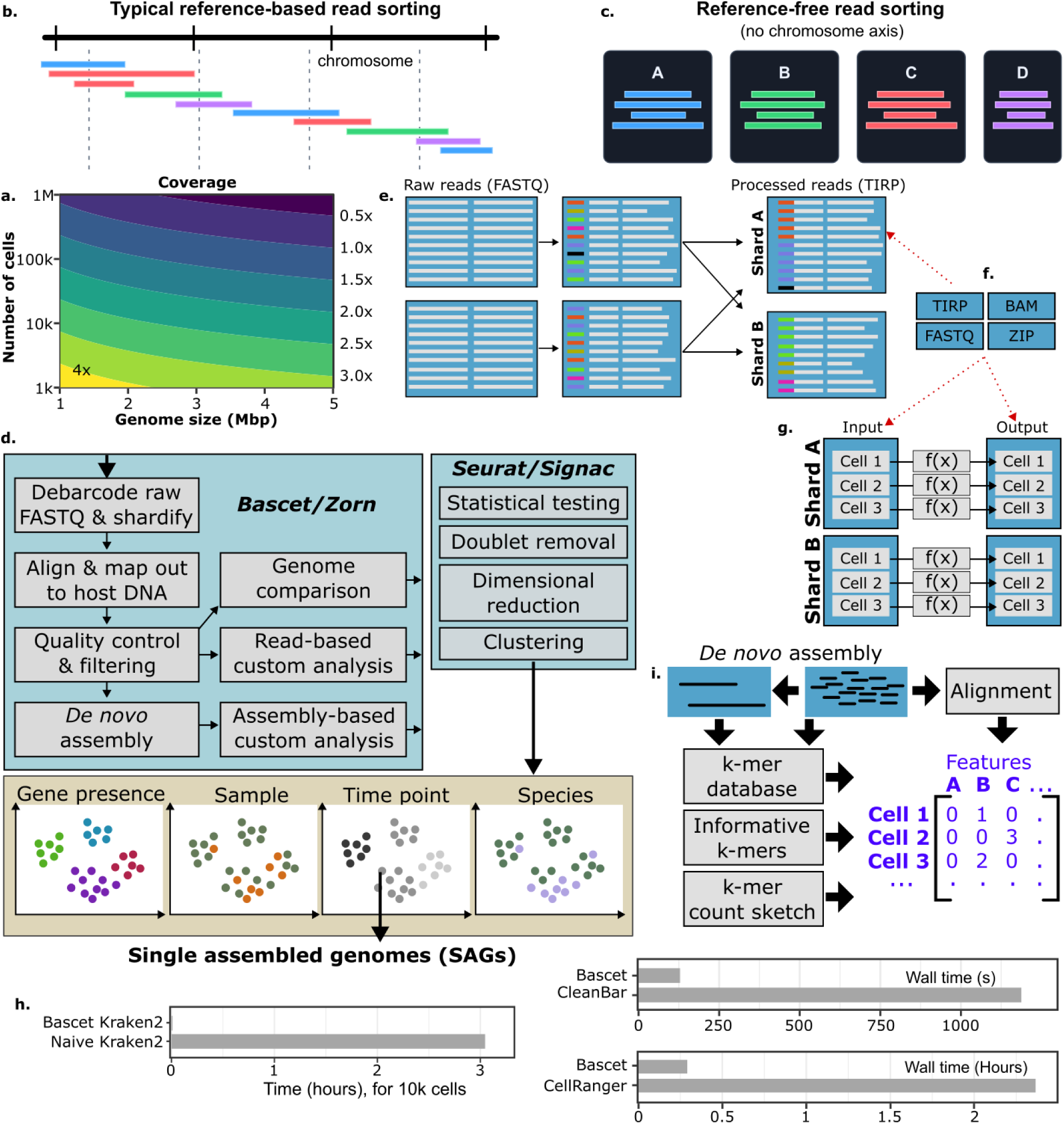
Bascet and Zorn enable large-scale scMetaG analysis. **(a)** Expected average coverage as a function of the number of cells and genome size, filling a Novaseq X 25B flow cell; coverage=25B/(GenomeSize*NumberOfCells). **(b)** Ordering of reads for typical single-cell RNA-seq (scRNAseq) analysis. **(c)** Ordering of reads for reference-free analysis, as supported by Bascet. **(d)** Typical overall scMetaG workflow, where the heavy preprocessing is performed by Bascet/Zorn, and most high-level processing by Seurat/Signac. The output is typically clustered and dimensionally reduced cells, where metadata and specialized analysis of each cell can be plotted to draw high-level conclusions. **(e)** Reads are processed and sorted into separate shards for fast parallel processing. **(f)** Bascet introduces several interchangeable file formats under a common API. **(g)** Bascet/Zorn support a general map-reduce procedure for applying a function (calculation) to each cell, both functioning as a minibatching workflow manager, as well as avoiding the creation of large numbers of files. **(h)** Benchmarks against CleanBar (a scMetaG debarcoding tool), Kraken2 (executed naively, on a cell-by-cell basis, versus Bascet’s implementation), and Cell Ranger (a popular commercial scRNAseq pipeline from 10x Genomics; see the **Supplemental Note** for details)**. (i)** To integrate with popular single-cell analysis tools, Bascet/Zorn generate features out of reads or contigs. Features are aggregated in a count matrix, handled similar to scRNAseq or ATACseq data.

To overcome these challenges, we developed Bascet, a highly scalable, open-source single-cell analysis pipeline, written in Rust. Conceptually similar to Cell Ranger (10x Genomics), Bascet is designed to perform common single-cell analysis tasks (e.g., barcode identification, read trimming/filtering, QC). However, unlike Cell Ranger, Bascet is able to perform reference-free analysis (**Fig 1bc, Supplemental Note**) and is agnostic to sequencing platform/chemistry, allowing it to accommodate diverse scMetaG protocols and the increasing number of new single-cell products coming to market. Further, additional scMetaG-specific methods are incorporated into Bascet (for e.g., taxonomic classification, *de novo* assembly, genome annotation), including novel methods for dimensionality reduction and data visualization (described below; **Fig 1d**).

To provide users with a familiar interface, we developed Zorn, a companion R package (**Fig 1d**). In addition to providing an interface between Bascet and popular single-cell analysis methods in R (detailed below), Zorn is a full workflow manager, inspired by Nextflow,^12^ enabling reproducible analysis. Because Bascet and Zorn are designed to be used together, we will refer to their combination hereafter as “Bascet/Zorn” when discussing methods that involve both tools.

Bascet/Zorn utilize several state-of-the-art strategies to enable efficient scMetaG analysis (**Fig 1efg**). First, we developed a novel family of file formats, all with a similar structure and unified under the common Bascet API (see Methods; **Fig 1ef**). Together, Bascet-formatted files limit data storage to a small number of files, thereby reducing small I/O operations and massively improving compute performance.

Building upon these novel file formats, we have further implemented several strategies to enable efficient computation (**Fig 1g**). Unlike typical scRNAseq protocols, most operations in the prokaryotic WGS space (e.g., *de novo* assembly, genome annotation) are performed on a “genome-by-genome” basis, in which sequencing reads, contigs, or scaffolds from a single prokaryotic isolate (or here, cell) are queried individually, one at a time. To enable analogous “cell-by-cell” operations here, we utilized the map-reduce concept from large-scale computing and functional programming (**Fig 1g**). Specifically, the Bascet map-operation applies a given function (a shell script or Rust function) over each cell, hiding how the data is actually stored from the user. This enables future addition of specialized file formats for increased performance and compression.

Finally, to maximize computational efficiency and enable scalable computing, Bascet/Zorn borrow the concept of “sharding” (horizontal partitioning) from databases (**Fig 1g**). Within this framework, data for a given cell is confined to a single “shard”. With each shard holding data for 1–10k cells, millions of cells can be managed without creating too many files for the filesystem to manage. This also reduces scheduling overhead for cluster job managers (e.g., SLURM), as each shard is a natural minibatch, enabling efficient multinode computing.

To showcase its efficiency, we benchmarked Bascet/Zorn against e.g., an existing scMetaG analysis tool (CleanBar, a scMetaG debarcoding tool),^13^ Kraken2 (executed naively, on a cell-by-cell basis, versus Bascet’s implementation),^14^ and Cell Ranger (a commercial scRNAseq pipeline from 10x Genomics; **Fig 1h**). In all cases, Bascet is substantially faster (>9x, 800x, and 8x faster, respectively; **Supplemental Note**).

Altogether, Bascet/Zorn enable efficient, flexible scMetaG analysis via novel file formats (improving I/O performance), map-reduce operations (enabling scalable cell-by-cell operations), sharding (maximizing computational resource usage), and chemistry/sequencing platform-agnostic design (software “future-proofing”).

### Bascet and Zorn enable flexible microbial single-cell analysis and integrate with popular eukaryotic single-cell tools

Most users will interact with Bascet/Zorn via Zorn in R, with minimal need for the command-line. While most users will follow a standard workflow (**Fig 1d**), processing steps are modular and can be added/omitted as needed. Briefly, users supply raw reads (FASTQ) to Bascet/Zorn, which are debarcoded and trimmed. At this step, users pick a suitable number of shards depending on how many computers are available for parallel processing (**Fig 1e**). This is followed by optional reference genome alignment, enabling e.g., host DNA removal (**Fig 1d**).

After debarcoding and trimming, we recommend users perform several Bascet/Zorn QC and filtration steps, specifically taxonomic classification and doublet/contaminant removal (**Fig 1d**). To perform a rough taxonomic classification of their cells, users can employ either (a) an “alignment-based” classification step, in which reads are aligned to user-selected reference genomes via BWA-MEM2/STAR (if sample composition is known), or (b) a “*de novo*” classification step, in which reads are assigned to taxa via Kraken2 (if sample composition is unknown).^14^ Both taxonomic classification steps allow users to identify potential biases (e.g., due to poor lysis of some taxa, poor sequencing coverage, contamination). Following taxonomic classification, users can detect and remove doublets (droplets/SPCs with >1 cell, from droplet instrument overloading or cell aggregation) and/or cells contaminated with free-floating DNA, using existing R packages (e.g., scDblFinder).^15^ Low-quality libraries can then be omitted from further analysis (**Fig 1d**).

After QC and filtration, users may wish to cluster and visualize cells/samples (e.g., via UMAP).^16^ To construct features for clustering analysis, users can choose from four workflows: (i) *alignment*, (ii) *k-mer database*, (iii) *informative k-mer*, and/or (iv) *k-mer count sketch* (**Fig 1i**). The (i) *alignment* workflow allows users to map reads and/or call variants relative to user-specified reference genome(s) via BWA-MEM2 and CellSNP-lite.^17^ The resulting read counts and/or SNPs serve as features for clustering and UMAP construction. Because the *alignment* workflow relies on reference-based mapping approaches, we anticipate that it will be most useful for comparing highly similar cells (e.g., members of the same species). For users interested in comparing more dissimilar cells (e.g., members of different species) or samples of unknown composition, we developed three *k*-mer-based workflows: (ii) a Kraken2-based *k-mer database* workflow, (iii) the MinHash^18^-inspired *informative k-mer* workflow, and (iv) a novel *k-mer count sketch* workflow (each detailed in subsequent sections; **Fig 1i**). All workflows enable UMAP construction and related clustering analyses, as Bascet/Zorn interoperate seamlessly with popular single-cell tools (e.g., Seurat, Signac).^19,20^

In addition to read-based analyses, users may also assemble SAGs *de novo* using SPAdes^21^ or SKESA^22^ (**Fig 1d**). Users may additionally co-assemble similar cells (defined by e.g., UMAP clustering) to improve coverage. In this scenario, Bascet/Zorn can be used to output reads of chosen cells for bespoke analysis.

Users may additionally run popular tools for microbial (meta)genomic data analysis, which are integrated into Bascet/Zorn (for e.g., read/SAG QC, genome annotation; **Fig 1d**). Results from these tools can be overlaid onto a UMAP (**Fig 1d**), alongside other metadata (e.g., taxonomic metadata, user-provided sample metadata), allowing users to interpret and explore their dataset.

Finally, the Bascet/Zorn *map* mechanism (**Fig 1g**) allows developers to add additional methods/tools. This feature will promote collaboration between members of the nascent scMetaG community and foster community-driven tool development. Altogether, Bascet/Zorn enable reproducible scMetaG data analysis, from start to end.

### Primary template-directed amplification in semi-permeable capsules enables high-throughput, high-coverage scMetaG

To showcase their utility, we used Bascet/Zorn to optimize a semi-permeable capsule (SPC)-based scMetaG method developed by Atrandi Biosciences, using an ATCC 10-strain (equal abundance) mock community (**Fig 2a-d**).^3,23^ Our scMetaG protocol consists of 6 major steps (**Fig 2a**): (a) encapsulating cells in SPCs, diluted to ensure singlets according to a Poisson distribution; (b) lysis; (c) whole-genome amplification (WGA); (d) debranching; (e) combinatorial barcoding of dsDNA using split-pool ligation;^24^ (f) barcoded DNA release, fragmentation, and index PCR.

**Figure 2.**
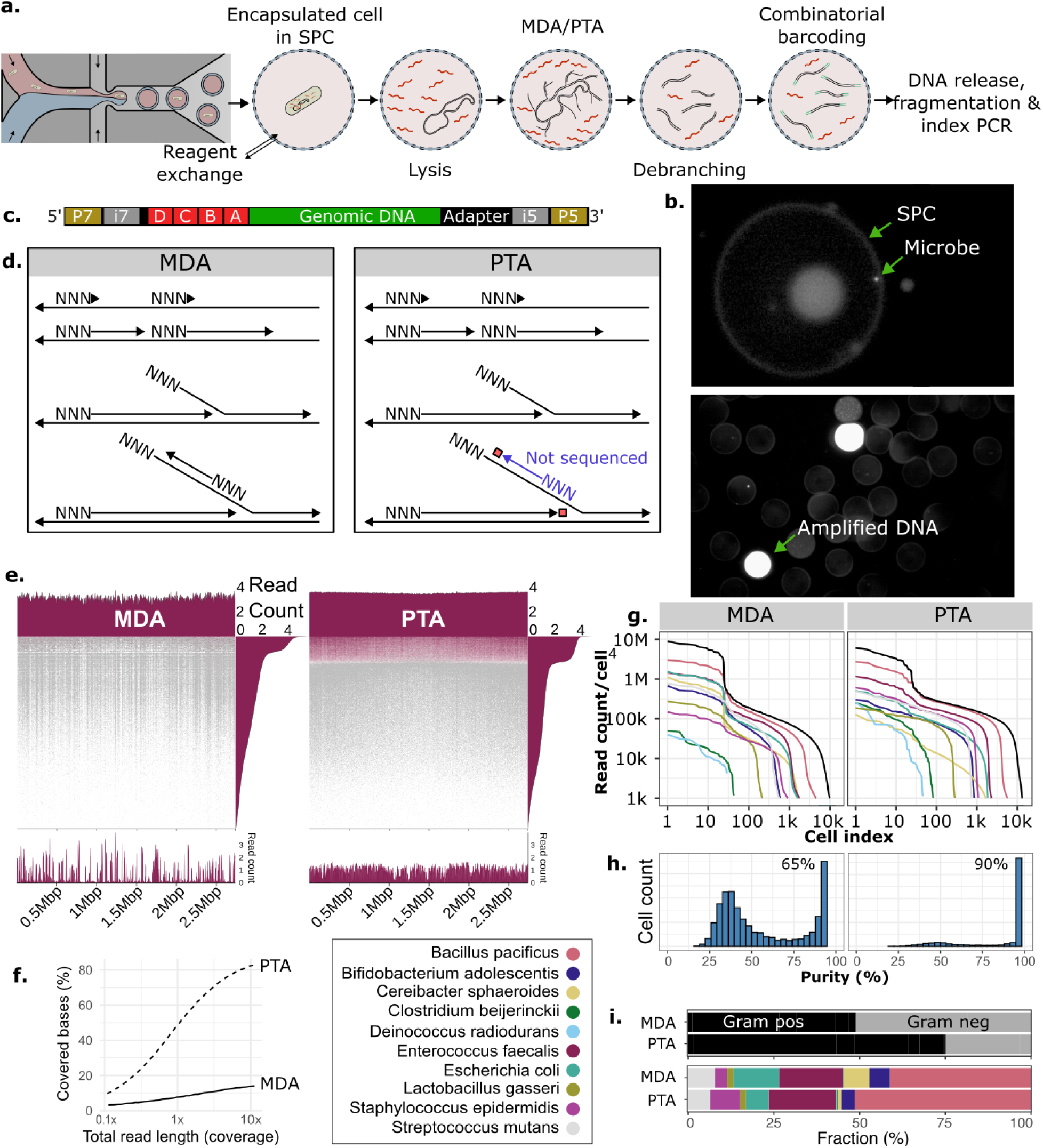
Semi-permeable capsules (SPCs) enable droplet-scale scMetaG of complex samples. **(a)** Our scMetaG protocol, adapted from Atrandi. SPCs allow buffers, small proteins, and small DNA oligos to be exchanged. After harsh lysis, genomic DNA is isothermally amplified using Multiple Displacement Amplification (MDA) or Primary Template-directed Amplification (PTA). After debranching to create simple double-stranded DNA (dsDNA), DNA is combinatorially barcoded. The SPCs are opened to release DNA, which is then fragmented to suitable size for sequencing. A final adapter is added, and after PCR, the library is sequenced. **(b)** Confirmation of cells in SPCs and Poisson loading, using microscopy (SYTO9 staining). Above: Encapsulated bacteria in SPCs. Below: stained DNA after MDA. **(c)** Final barcoded dsDNA, ready for sequencing. **(d)** Comparison of MDA and PTA protocols. PTA improves upon MDA by adding a blocking dNTP, preventing cyclic amplification, resulting in more amplicons of input template origin. **(e)** Coverage per individual cell genome (all species in **Supplemental Fig S1**). The number of reads is shown to the right. Below, local coverage for two cells having the same number of reads. **(f)** Library complexity indicated by coverage versus number of reads. **(g)** Knee plot, showing read count per SPC for MDA versus PTA, for each species (lysis protocol R4). **(h)** Histogram of fraction of reads assigned to the dominant species of a cell, i.e., library purity. Number of cells with >90% purity is indicated. **(i)** Relative abundances of species (based on read count) according to BWA-MEM2.

For (b) lysis, we benchmarked several protocols (see Methods), and derived a final protocol capable of capturing all strains. After lysis, we proceed with (cd) WGA and debranching. This generates overlapping amplicons, and unlike protocols that merely tagment DNA,^25^ enables *de novo* assembly of SAGs. Here, we benchmarked two WGA methods: multiple displacement amplification (MDA), which, despite being the most popular WGA method, is known to amplify unevenly;^21^ and primary template-directed amplification (PTA), the state-of-the-art WGA method in the eukaryotic scWGS space, which uses blocking dNTPs for more even amplification (**Fig 2d**). Crucially, we found that MDA library complexity saturates at 1x coverage, indicating that low complexity cannot be overcome by deep sequencing. Comparatively, PTA achieved more even genomic coverage (**Fig 2ef, Supplemental Fig S1**). We thus strongly recommend against MDA for scMetaG, and instead urge users to adopt PTA, as the latter greatly improves scMetaG data quality.

To assess mock community composition, we used Bascet/Zorn’s *alignment* workflow to align reads to mock community reference genomes. From a standard knee plot analysis, using total read counts per SPC, we identify 5k-10k cell-containing SPCs per sample (**Fig 2g**). However, knee plots for individual species indicate strain-specific lysis patterns, which are not captured in a standard knee plot (**Fig 2g**). We thus recommend low read count cutoffs and filtering cells after initial clustering. Using majority-class species assignments for SPCs, after doublet removal, <10% of PTA reads were contamination from other cells (**Fig 2h**). Further, our final lysis protocol was able to detect DNA from all 10 species (**Fig 2i**) and outperformed commercial protocols (**Supplemental Note**). Overall, SPCs in combination with PTA enable scalable, flexible, high-quality scMetaG protocols.

### Zorn’s *k*-mer database workflow leverages species assignments for clustering

Beyond the alignment workflow, we developed three *k*-mer-based workflows (*k-mer database, informative k-mer, k-mer count sketch*) for samples of unknown composition. We then applied each *k*-mer-based workflow to our mock community scMetaG data and compared them to the alignment workflow. The first *k*-mer-based workflow, Bascet/Zorn’s *k-mer database* workflow, uses Kraken2^14^ to obtain a table of read counts for each taxonomyID and cell, which can then be used for comparison. The quality of the resulting UMAP was comparable to that of the alignment workflow, despite Kraken2 assigning reads to taxa that were not present in the mock community (**Fig 3a**). Altogether, the speed of the *k*-mer database workflow makes it particularly suitable for initial QC and data reduction.

**Figure 3.**
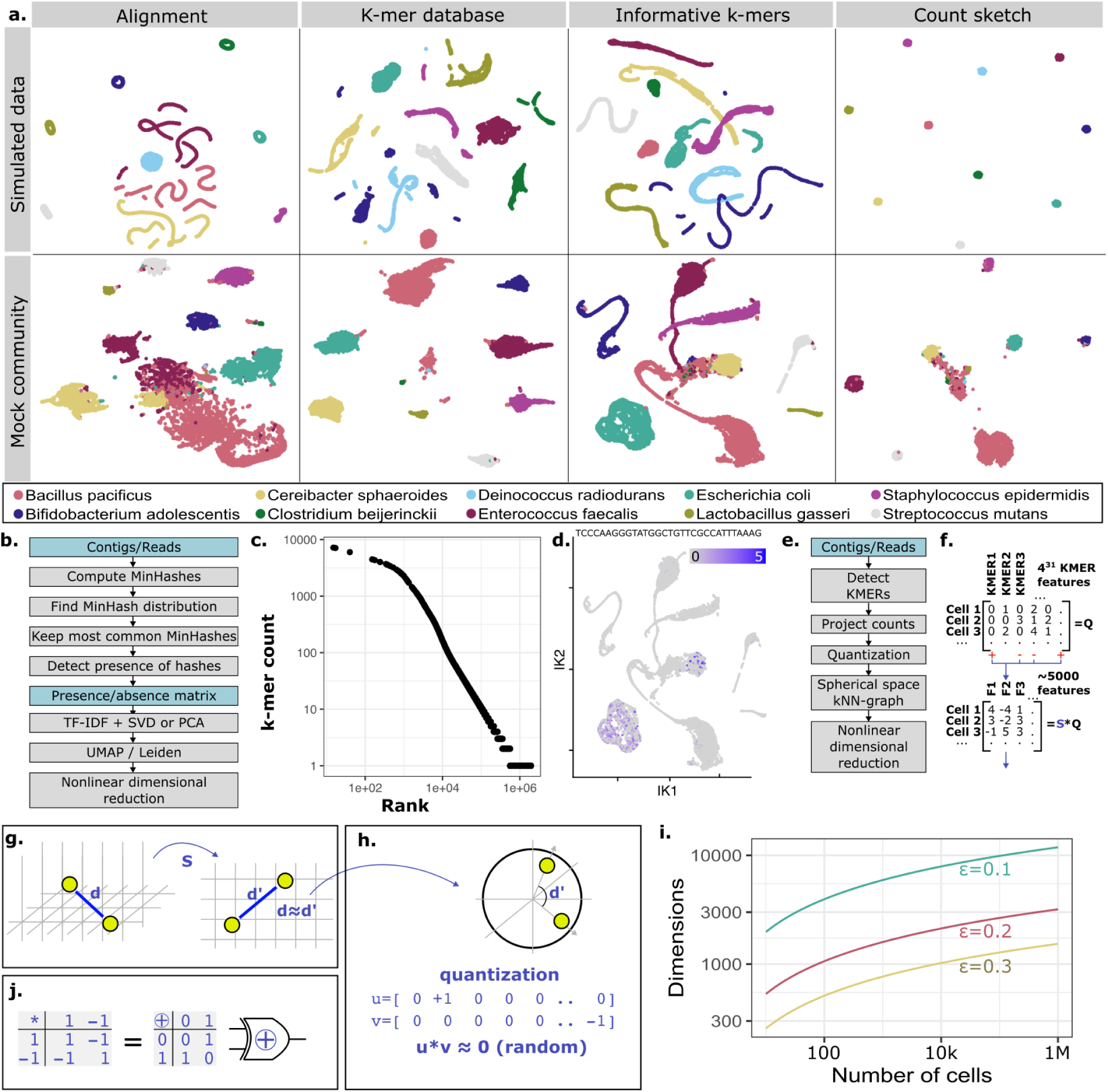
Bascet/Zorn workflow comparison. **(a)** UMAP of simulated and mock Primary Template-directed Amplification (PTA) communities, based on Bacet/Zorn’s four feature construction workflows. **(b)** The informative k-mer workflow. **(c)** Histogram of min-hash k-mer frequency. **(d)** Count of k-mer CAGATAGGGACCGAACTGTCTCACGACGTTC on UMAP. **(e)** The count sketch algorithm, including quantization steps. **(f)** Feature matrix, before and after count sketch projection. **(g)** Count sketch reduction randomly adds/subtracts k-mer counts from random columns in the final count matrix. This linear projection, which approximately conserves distances, can be performed without storing the vast amount of individual k-mer counts in memory. Features are then further decomposed to account for sequencing depth differences. **(h)** Cosine distances are distributed around zero for random zero-mean vectors. Binarizing vectors has little impact on angle. **(i)** Number of dimensions needed to conserve cell-cell distance with less than distortion ε. **(j)** Dot products of binary vectors are homomorphic to XOR functions, which are fundamental circuits, enabling the design of efficient computing hardware.

### Informative *k*-mers enable scalable and interpretable *de novo* comparison

Bascet/Zorn’s *alignment* and *k-mer database* workflows rely on reference genome(s) and a database of known taxa, respectively. As such, species/strains that are not represented in the reference set/database cannot be detected. We thus developed the *informative k-mer* workflow (**Fig 3b**), which uses a subset of *k*-mers detected across cells as features (**Fig 3cd**). *k*-mer subsets are selected (i) from the most common MinHashed *k*-mers, or (ii) by users (e.g., from genes of interest). Users can then proceed with e.g., UMAP construction, or use *k*-mer frequencies as input for Seurat differential abundance analysis.

To select an adequate *k*-mer frequency range, we tested an array of ranges and constructed UMAPs via the Seurat workflows for both ATACseq^26^ (TF-IDF, SVD, UMAP) and RNAseq (size-factor normalization, PCA, UMAP). We found that using the 100k most common min-*k*-mers produced a UMAP that was similar to those produced via Bascet/Zorn’s alignment and *k*-mer database workflows (**Fig 3a**).

Bascet/Zorn’s informative *k*-mer workflow enables interpretable *de novo* scMetaG analysis, without the need for reference genomes/databases. We anticipate this workflow will be particularly useful for users who lack prior knowledge of their samples, who prioritize the use of interpretable features (e.g., to assess which *k*-mers drive clustering; to identify differentially abundant *k*-mers via Seurat).

### *k*-mer count sketching leverages all *k*-mers for scalable *de novo* comparison

Because each *k*-mer sequence is known, Bascet/Zorn’s *informative k-mer* workflow enables interpretable *de novo* scMetaG analysis. However, the workflow is limited by the number of *k*-mers that can fit into memory. To enable use of the full *k*-mer composition space, we introduce a novel technique, namely a randomized affine projection to a subspace, implemented in Bascet/Zorn’s *k-mer count sketch* workflow (**Fig 3e-j**). This method is analogous to *count sketches*, which enable fast computation of the *k*-mer count projection, without intermediate storage of the full *k*-mer count matrix (which, for 1 million cells and 10 species, would occupy up to 2 petabytes). Bascet computes an initial randomized projection down to 5k dimensions (default), which Zorn can further dynamically reduce to trade precision for speed. The Johnson–Lindenstrauss lemma^27^ tells us that 5k dimensions are enough to roughly retain cell-cell distances (within 20%) for up to 10^6^ cells (**Fig 3i**). Since this is still a large amount of data for comparison, we sought fast approximations. We found that binarizing vector components to be ±1 both improved performance and data reduction (**Fig 3h**). A plausible explanation is as follows: (i) in high-dimensional spaces, randomly chosen vectors are highly likely to be orthogonal, and discretization affects the angle minimally; (ii) discretization also normalizes vector length, in a manner that appears numerically more robust than naive vector normalization. The discretized genome vectors are then compared using a fast approximate nearest neighbour algorithm, accessed via Seurat.

When applied to simulated and mock community data, the *k*-mer count sketch workflow produced exceptionally well-separated clusters, without prior knowledge of species or choice of *k*-mers (**Fig 3a**). We anticipate that this workflow will be useful for users with minimal prior knowledge of their sample and those interested in novelty detection/discovery.

### scMetaG provides maximum-resolution insights into the human oral microbiome

To test our full experimental and computational workflow on a complex sample, we used scMetaG and Bascet/Zorn to characterize saliva from a healthy human donor, generating 15,075 SAGs (**Fig 4ab**). When compared to bulk metagenomic data generated from the same saliva sample (see Methods), scMetaG generated substantially fewer human sequencing reads, and a larger proportion of reads from Gram-negative phyla (e.g., Bacteroidota; **Fig 4cd**).

Notably, scMetaG identified a small population of cells assigned to *Tannerella* sp. oral taxon 808 (TOT808), an uncultured, periodontitis-associated member of Bacteroidota,^28,29^ which was not detected using bulk metagenomics (**Fig 4e, Supplemental Fig S2**). When compared to the few available TOT808 genomes, a co-assembled SAG from our saliva sample represented a novel lineage within TOT808 (<97.3 average nucleotide identity with all TOT808 genomes; **Fig 4fg**). Interestingly, two major TOT808 clades consisted largely of genomes from either Asia or North America; our novel SAG was a member of the largely Asian clade (**Fig 4fg, Supplemental Fig S3-S4**). Overall, our results indicate that TOT808 are potentially geographically associated and further showcase the potential of Bascet/Zorn for large-scale SAG generation, inclusive of uncultured, low-abundance taxa.

**Figure 4.**
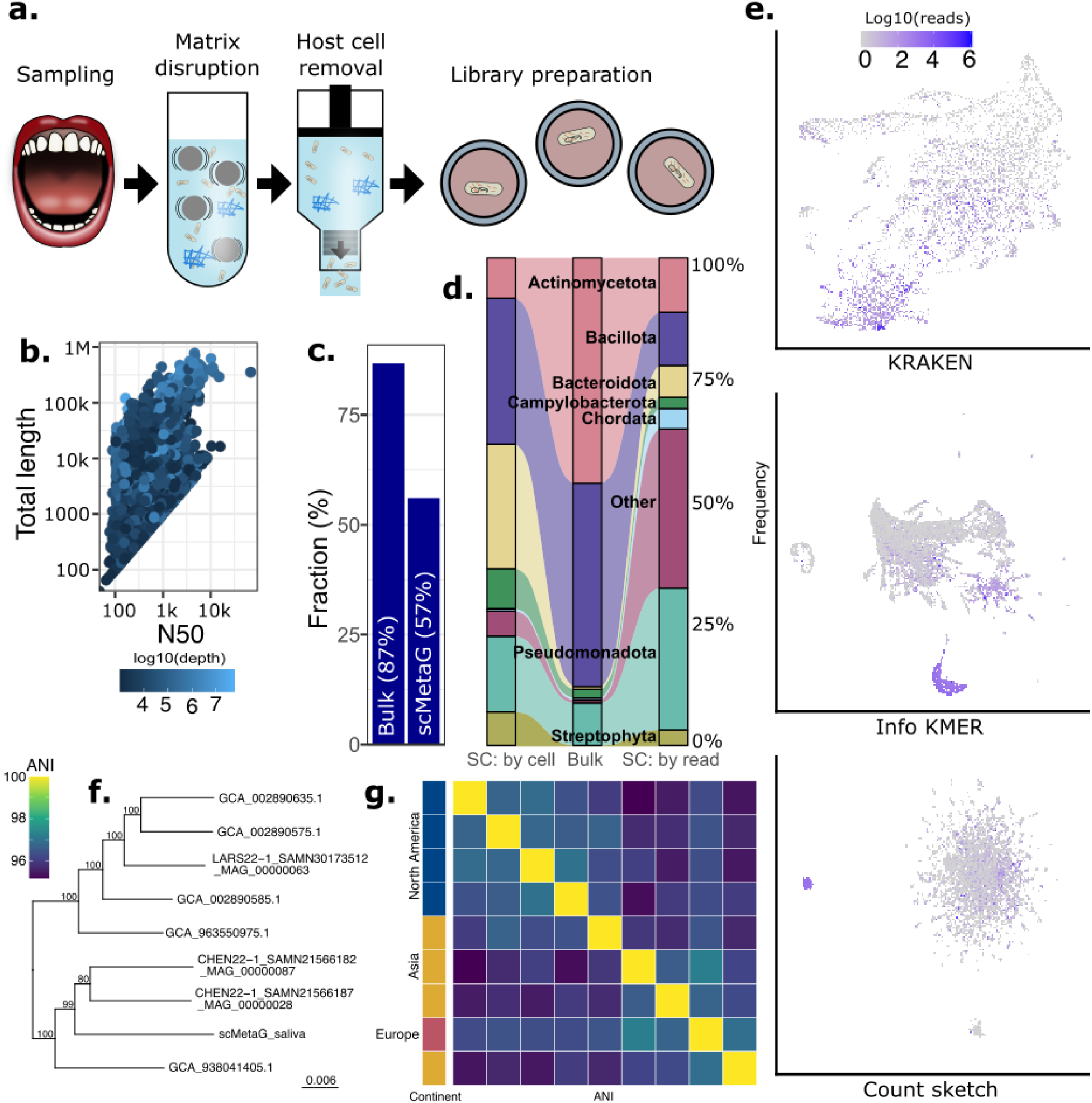
Single-cell-resolution genomics of 15k cells from the human saliva microbiome. **(a)** Saliva sample processing, prior to loading in semi-permeable capsules (SPCs). **(b)** Single-ampified genome (SAG) length (Y-axis) and N50 (X-axis), as a function of single-cell sequencing depth (color). **(c)** Percentage (%) of bulk metagenomic (left) and scMetaG (right) sequencing reads, which map to the human genome. **(d)** Relative abundance of phyla detected in saliva using (left-to-right): scMetaG, calculated at the cell level (“SC: by cell”); classic bulk shotgun metagenomics, calculated at the read-level (“Bulk”); and scMetaG, calculated at the read-level (“SC: by read”). **(e)** Human saliva UMAPs, colored by number of reads (log10[x+1]) mapped to Tannerella sp. oral taxon 808 (TOT808; NCBI Assembly accession GCA_002890575.1), produced using Bascet/Zorn’s (top-to-bottom) k-mer database (“KRAKEN”), informative k-mer (“Info KMER”), and/or k-mer count sketch (“Count sketch”) workflows. **(f)** Midpoint-rooted maximum likelihood phylogeny constructed using all high-quality TOT808 metagenome-assembled genomes (MAGs)/SAGs, plus a novel TOT808 SAG generated here (tip label scMetaG_saliva). Branch lengths are reported in substitutions/site. Branch labels denote IQ-TREE ultrafast bootstrap support percentages. **(g)** Heatmap of average nucleotide identity (ANI) values calculated between 9 TOT808 MAGs/SAGs.

## DISCUSSION

Single-cell (multi-)omic methods have revolutionized human biology. However, the single-cell ‘omics revolution has all but evaded microbiology, with only a handful of high-throughput methods in existence, including protocols for bacterial scRNAseq^30–33^ and, more recently, scMetaG.^3,6^ Single-cell sequencing technologies have the potential to usher in a new era of microbiology research, and with that comes new challenges and opportunities. On the experimental side, we developed a high-throughput scMetaG method using novel microfluidics that enables aggressive lysis and whole-genome amplification. We show that PTA, a state-of-the-art whole-genome amplification method, overcomes MDA’s severe amplification biases. Even when accounting for the increased cost of PTA, our scMetaG method is overwhelmingly more cost efficient than multiwell plate-based scMetaG methods (>150x cheaper, 0.27 EUR/cell; **Supplemental Note**).

While our optimal lysis protocol captures every microbe in a mock community and outperforms commercial protocols, lysis biases are still present. Further, our efforts centered around optimizing scMetaG for two types of samples (mock community and human saliva). In the bulk metagenomics space, microbiologists have spent decades developing and optimizing lysis/DNA extraction protocols for specific matrices/sample types,^34–39^ and we anticipate that the scMetaG community will benefit from similar endeavors. Specifically, novel lysis methods optimized for scMetaG droplets/SPCs are needed, as mechanical lysis methods, which work well for bulk metaG (e.g., bead beating)^40^ cannot be used.

Beyond high-throughput data generation, a lack of bioinformatic software for analyzing, storing, and disseminating microbial single-cell data further prevents microbiologists from making the single-cell leap. We provide Bascet/Zorn, the first bioinformatic toolkit for end-to-end analysis of large-scale microbial single-cell data, including novel file formats for efficient storage and data sharing. Unlike existing tools for single-cell analysis, Bascet/Zorn is designed to compare cells without a common reference. To handle the sparsity of single-cell data, we developed three novel *k*-mer-based methods. Most notably, we implement an approach based on randomized projections, utilizing all *k*-mers in the input data (the *k-mer count sketch* workflow). Interestingly, binarizing projected vectors improves species-species separation. Highly quantised representations have recently gained traction in deep learning, given their space efficiency and suitability for fast dedicated hardware (e.g., XOR operations; **Fig 3j**).^41^ Our approach translates cell comparison into a familiar problem within the field of machine learning, and is analogous to how alignment can be bypassed for the analysis of RNAseq data;^42^ our approach, however, is reference-free. This suggests that further theoretic unification of *k-*mer analysis may be fruitful. We anticipate that additional single-cell analysis methods can be incorporated into the Bascet/Zorn toolkit as they are developed, including additional QC steps, improved SAG (co-)assembly strategies, and incorporation of additional ‘omics layers (e.g., single-cell metatranscriptomics).

Overall, our novel scMetaG method queries microbiomes at maximum possible resolution (single cells) and boasts unprecedented throughput. Notably, we have the capacity to generate roughly as many microbial genomes in a single experiment as there currently are in existence in public databases. In the future, as data generation costs decrease, we anticipate that our scMetaG method and analysis toolkit will be particularly useful within the realm of diagnostics, pathogen surveillance, and precision health,^43–46^ likely in combination with cost-effective bulk methods. For example, researchers interested in linking plasmids to their host cells could combine scMetaG with long-read bulk metagenomics, the latter method providing more complete plasmids, and the former linking those plasmids to their host species/strain/cell. Likewise, researchers working with large numbers of samples could use shallow short-read bulk metagenomics to select a subset of samples for scMetaG. Altogether, recent developments here and elsewhere^3,30–33^ indicate that microbiology is on the cusp of a single-cell revolution.

## ACKNOWLEDGMENTS

This work was supported by: the SciLifeLab & Wallenberg Data Driven Life Science Program (grant KAW 2020.0239 to LMC); the Swedish Research Council (VR; grant 2023-05212 to LMC; grant 2024-03952 to JH; grant 2024-06085 to JH/LMC/JN; grant 2019-00453 to N Strömberg); the Swedish Foundation for Strategic Research (SSF; grant FFL24-0318 to LMC; grant ITM24-0035 to JH); Kempestiftelserna (IceLab Multidisciplinary Postdoctoral Fellowship Grant CSMK23-0129.3 to LMC/JH/N Strömberg; Kempe Postdoctoral Fellowship Grant JCK-0055 to JH); Cancerfonden (grant #23 3102 Pj to JH); the Umeå University Faculty of Medicine (grant FS 506-21 to JH/TL); funding from Institut Mérieux (the International Association for Food Protection [IAFP] and Institut Merieux Young Investigator Award in Antimicrobial Resistance to LMC); TUA #RV-977985 to N Strömberg. Computation was enabled by resources provided by (i) High Performance Computing Center North (HPC2N; Umeå University, Umeå, Sweden), and (ii) the National Academic Infrastructure for Supercomputing in Sweden (NAISS), partially funded by the Swedish Research Council through grant agreement no. 2022-06725. Experimental work was enabled by the μNisch Single-Cell Facility (Umeå University, Umeå, Sweden); the Umeå University Flow Cytometry Facility (Umeå, Sweden); Genomic Medicine Sweden Center North/Clinical Genomics Umeå, funded by a regional agreement between Umeå University and Region Västerbotten (ALF), Kempestiftelserna, SciLifeLab, Umeå University and Genomic Medicine Center (GMC) North; the National Genomics Infrastructure (NGI), funded by VR and SciLifeLab, Sweden.

## AUTHOR CONTRIBUTIONS

Software/computational method development was performed by HG, JD, AS, LMC, and JH, with input from TL. Computational analyses were performed by JD, JH, LMC, CS, and CD. Experimental method development was performed by IY, JV, and FA, with input from LM and JH. Clinical samples were provided by JN, N Sheng, and N Strömberg. LMC and JH conceived the study, which was funded by LMC, JH, TL, JN, and N Strömberg. All authors contributed to manuscript preparation.

## Online Methods

### Overall design of Bascet

To ensure speed and allow for the implementation of bespoke algorithms, Bascet was implemented in Rust (v1.93). All operations were implemented in a multithreaded fashion. To maximize inlining and reduce cache misses, the use of dynamic pointers was avoided. To enable future file format improvements, read and write operations are performed through one of the internal APIs that hide file format details. Bascet has several file formats including: Bascet-ZIP, a general-purpose format, which enables rapid storage/access to any set of files; Bascet-TIRP (Tabix-Indexed Read Pairs), a BED-like format for storing debarcoded sequencing reads; Bascet-FASTQ, designed to simplify interactions with tools that use FASTQ files as input (e.g., BWA-MEM2, Kraken2);^14,47^ and Bascet-BAM, a BAM/CRAM-like format^48^ that tracks cell origin. Early APIs were focused on reading one cell at a time, but this caused RAM usage spikes for large cells. The current APIs instead read chunks of reads at a time (sorted by cell ID), with downstream tools required to track which reads belong to which cell. As parallel processing primarily is over chunks and not cells, this only marginally complicates the workflow.

### Overall design of Zorn

Zorn was implemented in R (v4.4.0) to provide easy access to Seurat and Signac, with which it integrates, as well as to provide a familiar interface for bioinformaticians and biologists alike. The concept of shards was implemented in Zorn, where each file is stored as [name.##.ext]. The ## denotes the shard number, and each function call referring to “name” will automatically detect which shards belong together. Calls to Bascet give explicit paths to individual shards, meaning that Bascet need not be aware of the Shard concept. This implies that users are discouraged from using Bascet directly, and command-line invocations should rather be through R files and R scripts.

Computationally heavy Zorn function calls result in *Jobs*, which are executed using a *Runner*. There are presently three runners: (i) an immediate local runner, pausing R until the job is executed; (ii) an asynchronous local runner, returning immediately but expecting the user to check when the job is completed; (iii) a SLURM runner, which is inherently asynchronous. The local runner is only meant for developers of Bascet/Zorn, or new users who wish to test the workflow without committing to a full server farm deployment. Routine users are rather expected to set up SLURM and provide suitable parameters (e.g., account, number of cores), obtaining the full benefits of parallelization. To run all steps in a workflow, R should run on e.g., a login node using screen or tmux to keep the session alive. By default, all jobs are blocking calls (i.e., R is paused until the job has been executed). Optionally, a handle can be returned, which can be polled to see if the job is complete or not. This design is to benefit users who run RStudio via remote desktop (e.g., Open OnDemand) on the server farm, where one R session can be used to both quickly launch jobs while performing interactive data analysis.

Once Bascet has produced count data, Zorn can load it and provide it either as a Signac ChromatinAssay (if a reference genome is associated), or Seurat Assay for other workflows (e.g., *k*-mer or taxonomy counts). These objects can be combined into SeuratObjects to enable “multimodal” analysis of the same data, in the sense that different means of summarizing the cell data can easily be compared. The typical commands for e.g., plotting, differential expression, data integration, doublet detection, and imputation are then available.

### Detection of Atrandi WGS cell barcodes

Bascet has a modular system for handling different types of single-cell data. In the case of Atrandi WGS, the cells are first identified by a combination of four barcodes, each from a list of possible sequences (“whitelist”). This enables correction of the barcodes in case of sequence errors. The location of the barcodes within the read are first detected by analyzing the first 10k reads and scanning for whitelist entries. This step also prepares alternative positions to scan in case of a shift (e.g., ±1bp per barcode, and 4bp in total; tunable).

For performance, a naive algorithm first simply attempts to look up the barcode in the whitelist, from the expected position, using a hashmap. If this fails, the most similar whitelist entry is found by linear search. For performance, if any barcode fails, the algorithm stops. Barcode A is looked up first, as it is the most likely one to fail in case of over-fragmentation.

### Trimming of Atrandi WGS reads

Bascet performs initial trimming in parallel to barcode detection (see section “Detection of Atrandi WGS cell barcodes” above). Read 2 (R2) contains the cell barcode, and the initial portion is trivially removed based on the expected barcode size, plus 2bp to account for variation in dA-tailing. Read 1 (R1) is the genomic read, and must be trimmed if the insert is small and the read continues into the barcode. This analysis is complicated by the fact that there is no fixed adapter sequence. Initial use of fastp^49^ trimming (v0.23.4, default settings, paired mode) did not appear sufficient (some reads were still misclassified as human origin by Kraken2^14^ in the mock sample, and the countsketch was noisy due to residual barcodes). We thus implemented our own trimmer, executed in parallel with barcode trimming, as the barcode overlap in R1 and R2 can help trimming. A 16bp window is reverse-complemented and scanned for in the opposite read. Because the barcode is still present, this scan can proceed into it, giving high confidence in overlap for short reads. This ensures that residual barcodes are trimmed.

### Detection of Parse Biosciences and 10x Genomics cell barcodes

Bascet also supports debarcoding of raw reads from other single-cell chemistries. All currently existing versions of Parse Biosciences WT (whole transcriptome) are supported. As they are also combinatorial barcodes, they are detected analogously to Atrandi barcodes.

Barcodes for 10x Genomics RNAseq are also supported. As 10x Genomics has a much larger whitelist of barcodes, as opposed to combinatorial protocols, a different algorithm is used to find the most similar barcode. First, each first half of a barcode within the total set is stored into a hash set to deduplicate them. A vector of each unique first half in compact (16 bit) form is stored alongside a vector of vectors containing the full barcodes (32 bits) corresponding to each first half. Each barcode is looked up using a linear search within the much smaller vector of first halves. When a hamming distance between the searched-for barcode and the half is less than the current best match, the corresponding vector with full barcodes is searched for a match better than the current best. For 736,319 barcodes, the list of unique barcodes contains only 1,920, which amounts to only 3,840 bytes, which fits very well in level 1 cache. The secondary vectors each contain either 384 or 382 barcodes (4bp each) amounting to at most 1,536 bytes, which also fits in level 1 cache. SIMD is used for parallel hamming distance calculation.

### Correction of UMIs

Many single-cell RNAseq chemistries (e.g., from 10x Genomics and Parse Biosciences) include UMIs (unique molecular identifiers) to track the origin of cDNA and enable deduplication. Bascet implements an algorithm analogous to the UMI-tools directional algorithm.^50^ For single-cell WGS (including scMetaG), deduplication is not necessary, and the UMI field is simply set to be empty.

### Sharding and design of the TIRP file format

To make the remaining steps agnostic to the choice of library preparation method (i.e., to support other protocols than those described in this manuscript), the debarcoded and trimmed reads are stored in a new TIRP (Tabix-indexed read pairs) file format. This is a modified BGZIP-sorted BED file, enabling the use of Tabix^51^ to retrieve all reads for a given cell barcode. The file has 8 columns (chrom, from, to, r1, r2, q1, q2, UMI), where r# and q# represent read and quality score. The first three columns are mandated by the BED standard. By setting chrom=cellbarcode, from=1, to=1, and sorting the file, Tabix can extract reads for a cell by asking for all reads from the given “chromosome” (i.e., the name of the cell is used). While the from–to columns are redundant, compression of the file will largely remove them. For Atrandi scMetaG, UMI=””, as the field is primarily intended for scRNAseq protocols. Alternatives to TIRP were considered and may be implemented in the future, as the Bascet API enables easy addition of new file formats. TIRP, however, has the advantage that users can efficiently extract the reads for a particular cell using Tabix.

To maximize multiprocessing, each raw FASTQ file is debarcoded, trimmed, and outputted as a TIRP file that is sorted. As sequencing may be performed on multiple lanes, one library may result in multiple FASTQ files. To shard the files, these FASTQ files must be merged, and simultaneously rewritten, but split on a cell-by-cell basis. This merge has to be performed on a single compute node, making it a workflow bottleneck. By pre-sorting the TIRP-files, sharding becomes almost O(number of reads), making it primarily bound by disk I/O speed.

### High-performance sorting of reads per cell and TIRP file generation

The design of Bascet further requires data to be sorted by cell identity to limit memory usage. In sorted data, encountering a new cell identity guarantees that all data for the previous cell has been seen, allowing its results to be written to disk and freed from memory. Without sorting, results for all cells must be held simultaneously, as any cell may appear at any point in the file, and memory usage scales with the number of cells.

While utilities exist for sorting reads by position (e.g., SAMtools), there are no utilities designed to sort reads by their cell of origin (besides storing each cell in separate files, which is inefficient). Early Bascet versions used the Unix sort program on uncompressed BED files (TIRP files are BED files with specific columns), followed by compression (into BGZF, Blocked GNU Zip Format). However, this approach resulted in a large amount of disk space used, and several hours of processing. However, in-memory sorting is limited by available memory (RAM), while external sorting is I/O bound and operates on fully decompressed data, increasing the volume of data written to and read from disk. We figured that sorting already compressed data, and blocks of reads belonging to the same cell, would speed up the processes. However, BGZF files do not provide a way to determine which data belong to which cell without full decompression. BGZF is a commonly used bioinformatics file format built on the GZIP format^52^ that divides data into blocks that each support metadata through the GZIP extra subfields. This mechanism is never used for cell identity, remaining limited to the block-size (BSIZE) field;^53^ however, it is precisely what would make cell identity accessible at the block level without decompression. If cell identity were encoded in the block headers, sorted data would allow downstream tools to detect cell boundaries by reading the GZIP header alone, rather than decompressing every block. Block-level cell identity further provides index-like selective access without a separate index file at a very low computational cost.

To solve this problem and enable fast compression and sorting of TIRP files, we developed the BBGZ format (**Bascet block gzip; Supplemental Fig S5**), which enables an alternative workflow that avoids both bottlenecks, while remaining backwards compatible with existing BGZF tools (**Supplemental Fig S6**). During demultiplexing, reads are assigned cell identities and locally sorted in memory into 1-GiB sized chunks, each compressed as BBGZ. Because each BBGZ block contains only records matching a single cell identity, local sorting within a chunk reduces to grouping records by their ID subfield rather than sorting arbitrary data. These locally sorted chunks are then merged into a globally sorted output using a K-way external merge^54^ that operates exclusively on BBGZ block headers. Because cell identity is encoded in the ID subfield, the merge needs only to compare header metadata to determine block ordering, requiring no decompression at any stage. As sorting operates on compressed rather than decompressed data, memory and I/O scale with the compressed data volume. This reduced peak memory usage by >64-fold (4 GiB required against ∼256 GiB previously recommended) and runtime by >10-fold compared to the conventional workflow. Both improvements scale linearly with dataset size.

By moving cell identity labels to the block level, the BBGZ format enables high-performance, low-memory sequential access of data and index-free (retaining index-ability) random access. We thus reduce sorting time greatly (benchmarks in **Supplemental Note**).

### Design of the Bascet-ZIP file format

As we expect to wrap many existing tools, there is a need for a structured approach to handling their input and output. One approach is to force the user to standardize the output from each software; however, it is near impossible to find a one-size-fits-all solution (e.g., conversion to JSON or XML), and this would incur a heavy burden on the developers that wrap upstream software. We thus instead aim to store the raw output data as-is. The current exceptions are (i) contigs (FASTA-like data) and (ii) raw reads and quality scores (FASTQ-like data), as these files will commonly be piped across tools, calling for a standard format.

Several storage solutions were considered, and through the Bascet API, improved file formats can easily be added. The use of a separate object store (e.g., S3, MongoDB) was ruled out, as such solutions require the maintenance of separate persistent processes, and many compute facilities do not support them. Furthermore, for most of our envisioned users, object stores are difficult to copy between machines or make backups of. We instead opted for a single container file, aimed for (i) immutable data (data is only generated, never modified), (ii) fast listing of content, (iii) reading of files in arbitrary order. All of these requirements are met by basic ZIP files, as they contain an index to all files within. Experiments showed that writing a ZIP-file having 100k (trivial) files can be performed in seconds, while a regular filesystem would struggle to cope with this quantity of files (see **Supplemental Note**). As an example of the latter, remote copying using scp may take nearly a second to set up the transfer of a single file, no matter the size of the content. However, by keeping all small files together in a ZIP, only one such handshake is needed, and disk prefetch results in maximum I/O performance.

The current ZIP format is rather trivial, in that files are stored as #/randomfile.xyz where # is the cell barcode. For FASTQ-type content, the data is stored as #/r1.fq and r2.fq.

Contigs are stored as #/contig.fa. A special file, #/_mapcell.log, contains the stdout of the tool that was used to generate the content. Other tool-specific log files may also be stored, normally under their original names.

To enable random I/O to files within the ZIP-file, the files can be stored in uncompressed format. It is the responsibility of the storing process (e.g., the map-reduce system) to ensure that this happens. The Bascet API can then extract the start and end coordinates of the file within the ZIP-file, enabling advanced tools to directly look up needed data. An example use case is to store per-cell *k*-mer databases, and enable the lookup of individual *k*-mer counts without the need to first extract the entire *k*-mer database. This is thus crucial for the performance of some envisioned future tasks.

### Design of the map-reduce system

To apply functions to each cell’s data, we designed a system around constraints imposed by existing bioinformatic software and the file formats supported by such software. This has resulted in a general API, provided as several Rust traits, each available if the underlying file format enables the implementation.

Two modes of data reading are supported: (i) streaming, which reads all objects in the file, and (ii) random, which specifically reads requested objects. Since random access requires seeking based on an index, not all file formats support this (e.g., Bascet-FASTQ has no means of seeking to a particular cell). All file formats support streamed access. Random access is only beneficial in a few scenarios: (a) when extracting a filtered list of objects, (b) when extracting a single object, e.g., the assembled genome of a particular cell of interest.

The function to be applied is, on the highest level, a function in Rust. Functions written in this format have the highest level of possible performance, and can e.g., read specific parts of objects in the container without extracting the full file. This is useful when embedding per-cell databases in the container file. As this type of function requires knowledge of Rust, we expect only a handful of power users to make use of it.

The most commonly used Rust map function takes a Bash script as input and runs the script as the map function. This approach is designed to be suitable for a general user who wishes to wrap existing software into Bascet/Zorn. For this approach to work, the Bash script must provide a list of files that the software needs. The underlying API will then create these files based on how the container file format is designed; e.g., a requested r1.fq will simply be unzipped from a Bascet-ZIP, but created *de novo* if the input is a Bascet-BAM. The downside of this approach is that an additional file creation step is needed; however, not all software can pipe two read files, and this approach is more amenable to parallelized decompression. The Bash script also specifies the preferred number of threads; if this is more than one thread, priority is given to distribute as many threads as possible to a smaller number of map-scripts. This turns out to (i) reduce peak main memory usage, and (ii) solve a bottleneck where a map-reduce operation waits for a single cell with large amounts of data to be processed to completion.

The reduce step is performed using the Zorn *aggregate* function. Two key assumptions went into this design: (i) reduction is computationally cheap, within the capacity of a single computer; (ii) reductions may have rich structure, such as being a tree or R S4 class. Performing the reduction largely using R code fits these assumptions, where Zorn calls Bascet merely for extracting files from the container object.

An issue with the map-reduce system is that it relies on external software. Early versions of Bascet aimed to deliver all required software via Conda, wrapped in a Singularity or Docker container. This however ran into multiple issues: (i) Conda failed to resolve all versions of all upstream software once the number of software grew past a critical number; (ii) the time to start the container grew to near 5 seconds, making it too inefficient to call Bascet for each file to be extracted; (iii) different platforms required different containers (including Podman), making it too challenging to maintain all options. Routing absolute paths on different operating systems was difficult, especially for Windows. Of all these problems, we could work around the startup issue (ii) by creating a special Bascet mode enabling streaming of commands through a live connection. This connection was kept alive using the R processx library (v3.8.6). The solution is fragile due to the lack of R error handling and will likely be removed in future versions (likely by wrapping Bascet using R FFI).

To overcome the integration issues, we instead translated the upstream software to Rust, as described in separate work.^55^ The versions of translations are listed in the **Supplemental Note** (such software will here be denoted “*translated*”). This also resulted in speed improvements, as data could be sent directly to the relevant functions without reading and writing intermediate files. Future versions of the map-reduce system will likely assume that all software is translated, as this is a prerequisite for high performance single-cell analysis.

### *De novo* assembly

Bascet wraps both SPAdes^21^ via the MAP system, and SKESA^22^ (*translated*). Each *de novo* assembler outputs a FASTA file containing contigs. These FASTA files are stored in Bascet-ZIP as described above (see section “Design of the Bascet-ZIP file format”). SPAdes supports assembly of single-cell genomes, amplified using MDA, taking into account the local-exponential nature of the reaction (i.e., via its single-cell assembly mode; option ‘--sc’). However, we find that, for the massive number of libraries that our scMetaG method generates, single-cell SPAdes is too slow to be practical. We thus used SKESA v2.5.1 for large-scale analyses conducted here (e.g., assembly of >15k SAGs from saliva; ‘--min_contig 50 --min_count 2 --fraction 0.1’), and single-cell SPAdes for targeted analyses (e.g., co-assembly of the saliva TOT808 SAG; ‘--sc’).

### Alignment workflow

Bascet wraps several aligners, including BWA-MEM2^56^ (*translated*) and STAR^57^ (*translated*). These aligners cover the needs of DNA and RNAseq reference-based alignment, respectively. To keep the read-cell barcode linkage, a custom FASTQ file is written, where the name of the cell is encoded as the name of the read. This implies that certain characters are not allowed in cell names (e.g., whitespace). After the aligner has produced a BAM file, Bascet optionally sorts and indexes the file (used for counting of features, such as for RNAseq and ATACseq analysis). Reads are otherwise ordered such that all reads for one particular cell belong together. Bascet then provides the means of counting the reads mapping to a reference genome, storing the counts in an HDF5 file format using sparse encoding. As one count table is produced per shard, Zorn takes care of merging count tables upon loading. Within Bascet’s alignment workflow, users additionally have the option of performing variant calling using CellSNP-lite.^17^

### *k*-mer database workflow

Bascet wraps Kraken2^14^ (*translated*) for read-based taxonomic classification. The standard database is standard-8, as we envision that this workflow will primarily be used for rough classification, and thus the smallest database is the best fit. Because Kraken2 spends significant time loading the *k*-mer database, the reads for all cells have to be processed in a single invocation. This influenced the creation of Bascet-FASTQ and Bascet-BAM as intermediate formats to support such tools. In addition, Bascet can directly send reads to functions of programs that have been translated to Rust, bypassing additional writing and parsing. The original Kraken2 also generates a large uncompressed output file with each read classified. As a large amount of time is spent writing this file, it has been made optional in our translation. Instead, the primary output is a count matrix (sparse matrix anndata HDF5 file^58^) of taxonomyID *vs* cell name, that is produced directly, bypassing the need for re-parsing the Kraken2 classification file. After loading this matrix using Zorn, cells can be assigned a consensus taxonomy ID (i.e., assigned to the most common ID among reads for the cell). To aid visualization, Zorn can further map consensus taxonomy ID to the name of phylum, class, order, family, genus, and species (internally using taxonomizr v0.10.7; https://CRAN.R-project.org/package=taxonomizr).

### Informative *k*-mer workflow

Informative *k*-mers can be extracted both from raw reads and contigs produced via *de novo* assembly (see section “*De novo* assembly” above). As contigs cannot currently be produced for cells with extremely low coverage, we have prioritized the read-based workflow.

In early versions of Bascet, we first deduplicated *k*-mers using KMC3.^59^ We decided against this approach, as it was slow, and we encountered random crashes in the KMC3 ops-step (where we merged *k*-mers across cells). Furthermore, the count of *k*-mers may also be informative (e.g., possibly reflecting plasmid copy number). Thus, the current version counts *k*-mer (and *k*-mer hashes) directly from reads without any attempt at pre-sorting them. An efficient rolling hash algorithm was used, as it enabled fast computation for overlapping *k*-mers^60^ (nthash-rs v0.1.3).

After the user has selected *k*-mers of interest, all reads are scanned again. While looking up *k*-mers in a *k*-mer database may theoretically be faster, it is likely that the large number of queried *k*-mers will be randomly scattered over the database. Thus, due to cache misses and how the disk is read in blocks, any theoretical speed gain would likely not pan out in practice.

To select an adequate *k*-mer frequency range, we empirically tested *k*-mer frequency ranges and constructed UMAPs via the workflows for both Signac ATACseq^26^ (TF-IDF, SVD, UMAP) and Seurat RNAseq (size-factor normalization, PCA, UMAP). We initially expected the most abundant *k*-mers to be of low utility; however, we do not appear to find any such *k*-mers, most likely because the *k*-mer length (31bp) results in near species-specific *k*-mers. For the mock community, at our sequencing depth, at least 10k of the most common MinHash *k*-mers are needed to ensure that cells cluster together. However, with about 100k *k*-mers, we observe a drastic improvement in clustering, and thus used the top 100k most common MinHash *k*-mers in our analysis.

### *k*-mer count sketch workflow

We aim to be able to model the distribution of counts *X* for *k*-mer *k*, from the genome *z* of cell *i*, as

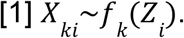

The latent space, *Z*, can be equipped with a normal prior to make this a typical generative process. We will however not observe *X* directly (unless we assemble the genome), but rather sampled *k*-mer counts, *Q_k_*. Ignoring the correlation between *k*-mers and local exponential amplification biases of multiple displacement amplification (MDA),^21^ a simple model is

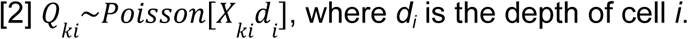

where *d_i_* is the depth of cell *i*.

In practice, we simply use the depth estimate

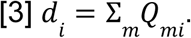

To compute a lower dimensional representation, *Z*, we decided to use UMAP.^16^ This algorithm takes as input the *k*-nearest neighbour (kNN) graph (i.e., most similar cells). Given above, we can compute this graph over the estimated *k*-mer distribution,

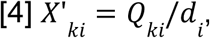

where *d_i_* takes the role of a size factor for cell *i*. As the distance between points, we use the cosine distance, as the probability of random vectors being orthogonal (high distance) increases with dimensionality. This has previously been shown to benefit single-cell analysis.^61^

Computing the kNN from Equation [4] is, however, intractable as *Q* has 4^31^ features (or about 5x10^7^ for a mix of 10 different bacterial species) and cannot be kept in memory (merely keeping the features sorted on disk is a challenge). We instead compute a randomized projection, *SQ*, to a lower-dimensional zero-centered space,^62^ where *SQ* largely keeps the structure of *Q*; e.g. ||*SQx*||_2_ = (1±ε)||*Qx*||_2_, and low rank approximations Q=LDW can be found (where if Q is n × n, L is n × k, D is diagonal k × k, and W is n × k). Such approximations also open up for future alternative dimensional reduction methods, as well as generative process formulations. In the explicit algorithm, which turns out equivalent to the count sketch algorithm,^63^ *S* is defined through hash functions h(x) → {1,…,n} and g(x) → {-1,+1} as

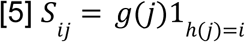

For h, the bucket index was derived from the lower 32 bits of a canonical ntHash2^64^ hash, masked against the sketch size minus one via bitwise AND. For g, the sign was derived from the same canonical ntHash2 value by rotating the hash right by 32 positions and reading the least significant bit, mapped to +1 or −1. Due to the linearity of the mapping, the entries SQ = S(Q1 + Q2 + …) can be computed in parallel using multiple threads. Furthermore, there is no need to ever retain *Q* in memory, meaning that the memory requirement is determined by the size of *SQ* only. To help fine-tune the size of the final projection, without recomputing SQ from all raw data, Zorn may further apply a second randomized zero-centered projection as *TSQ*. *T* is defined as

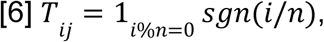

or equivalently, for each output vector, an equal number of input feature vectors are added and subtracted. *S* and *T* both transform the features such that they are zero-centered, motivating the choice of cosine distance.

Before computing the kNN-graph from *SQ*, we tested different methods of normalization, where division by *d* in Equation [4] is the natural approach. While the normalizing factor, *d*, can be computed from *SQ*, computing it with good precision appeared difficult (using standard double floating point), and the resulting UMAPs were of poor quality. Inspired by product quantization (which can be used to speed up computation while also reducing memory footprint)^65^ and binarized neural networks,^41^ we instead investigated the quantized

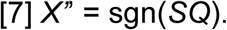

This method results in trivially normalized vectors as ||*X*’’ || = 1. For the cosine distance, it retains the key property of large vectors, namely their low probability of randomly being aligned. To finally compute the kNN graph, an approximation was needed, as computing all cell pairwise distances was prohibitively slow (even for 5k dense dimensions). We thus approximated the kNN graph using the library uwot.^66^

### Reference genome-based analysis of mock community data

A reference genome was constructed by concatenating the 10 reference genomes of the ATCC mock community strains (MSA-2003; genome sequences obtained from ATCC) with *Xanthomonas campestris* B-1459 (per the addition of xanthan gum in the scMetaG protocol; NCBI Assembly accession GCF_001372255.1) and phiX (Illumina). BWA-MEM2^56^ (*translated*) was used for alignment via Bascet/Zorn (default settings).

Reads that mapped to either genome were conservatively removed for raw data deposition and further analysis. To analyze the fraction of reads mapping to either reference genome, the CIGAR string was analyzed, and only reads were considered to be mapping if BAM unmapped flag (0x4) was clear and at least 90% of its read bases were mapped (CIGAR M).

To assess mixing cells, we compared the counts of reads for the dominant species for a cell *vs* the total count of reads of other species (as assigned via Kraken2 and the standard-8 database, or by alignment for the mock community).

### Generation of simulated mock community data

An R script was written to simulate reads from cells, immediately creating a TIRP file (i.e., the barcoding step was skipped, as this benchmark focused on assessing the performance of Bascet/Zorn’s dimensionality reduction methods). For genomes consisting of multiple contigs, an equal ratio was assumed. The MDA amplification and library preparation was assumed to be ideal, such that reads were simply uniformly and independently sampled across the genome of each cell. All cells were singlets. The insert size of all fragments was assumed to be 800 bp, resulting in 2x150bp reads. The number of reads per cell was sampled uniformly (log scale), from 10^3^ to 10^4.5^ reads. For each species, 50 cells were simulated.

NCBI accessions for simulated genomes were as follows: *Bacillus pacificus* (CP086328.1, CP086329.1), *Bifidobacterium adolescentis* (NC_008618.1), *Cereibacter sphaeroides* (NC_009049.1, NC_009050.1, NC_009040.1), *Clostridium beijerinckii* (CP006777.1), *Deinococcus radiodurans* (CP150840.1, CP150841.1, CP150843.1, CP150842.1), *Enterococcus faecalis* (NC_017316.1 and NZ_OQ434560.1), *Escherichia coli* (NZ_LR881938.1), *Lactobacillus gasseri* (NC_008530.1), *Staphylococcus epidermidis* (NC_004461.1), *Streptococcus mutans* (AE014133.2).

### Saliva sample collection and processing

Fresh saliva was obtained from a healthy (non-patient) human donor (ethical permit # 08-047M). The saliva (5mL) was collected in a 50mL falcon tube and vortexed for 3 min with glass beads (Fisherbrand, catalog #41401001). The sample was subsequently filtered using a 35μm filter (Cat #352235, Corning). The filtrate was further centrifuged at 5,000g for 10 minutes. The supernatant was discarded and the pellet was washed with 1mL of PBS three times. Finally, the pellet was resuspended in 10% glycerol and stored at -80°C.

### Single-cell bacterial encapsulation in semi-permeable capsules

Samples (i.e., ten-strain mock community ATCC MSA-2003 and the processed saliva sample; see section “Saliva sample collection and processing” above) were diluted before starting the single-cell encapsulation using phosphate-buffered saline (PBS). The dilution was made based on the cell count as observed by counting the cells stained with 10x SYTO9 nucleic acid dye (Cat #S34854, Invitrogen) on a hemocytometer to get λ of 0.1 or lower for final droplet loading.

The ONYX platform (Atrandi Biosciences, Cat #MHN-ONYX1) was used for SPC production using the SPCs innovator kit (Cat #CKN-G-12, Atrandi Biosciences). The procedure was followed as per the instructions in the manual for the preparation of the core and shell solutions used for SPC generation. The saliva sample was pretreated with DNase I (0.1U/ul, NEB, 37°C for 30 minutes) for the removal of any host or free-floating DNA. This was followed by the addition of 0.05% Xanthan gum (Cat #G1253, Sigma) to the final core solution, which helps to prevent any aggregation of bacteria together or to the channel walls, thus helping to maintain a smooth flow.^67^ The encapsulation was performed for 3h, and the excess oil was removed from the emulsion collected in the collection tube. The shell was crosslinked using a UV light source of 405nm (Atrandi Biosciences Light Exposure Device) for 30–60s, followed by emulsion breaking and washing the capsules with Capsule Wash Buffer (CWB). The occupancy of the capsules was further checked by fluorescence imaging using 10x SYTO9 nucleic acid dye.

### Lysis of cells

Several lysis protocols were tested, with the full list of protocols and sequencing data available in the **Supplemental Note**. For the datasets presented in the figures of this paper, we used lysis method R4, which is inspired by a previous study.^6^

Bacterial lysis protocol R4: The SPCs were fixed in chilled 100% methanol before the initiation of the lysis. SPCs were divided into different tubes and incubated in enzyme cocktail solution with TE buffer (pH 8) containing 5U Lysostaphin (Cat #L9043, Sigma), 50U Mutanolysin (Cat #SRE0007, Sigma), 50U Lysozyme (Cat #E-0057-D2, Biosearch Technologies), 0.5mg/ml Achromopeptidase (Cat #A3547, Sigma) with 2.5mM EDTA and 10mM NaCl at 37°C with shaking at 800rpm overnight. The next day, the SPCs were washed three times with WB and further resuspended in the Proteinase K lysis solution in TE buffer containing 4U Proteinase K (P8107AA, NEB), 1% Triton X (Cat #X100, Sigma) and 100mM NaCl. The incubation was done at 55°C for 1 hour with shaking at 800rpm. After incubation, SPCs were pelleted and resuspended in 1 mL of 4M guanidine thiocyanate (GTC) and incubated at room temperature for 30 min. The SPCs were spun down and washed three times with WB and two times with nuclease-free water.

### Multiple displacement amplification and cell labelling

Multiple Displacement Amplification (MDA) and cell labelling was carried out using the Single-Microbe DNA Barcoding Kit (Cat #CKP-BARK1, Atrandi Biosciences). Briefly, SPCs were mixed with all reaction components for WGA and incubated at 45°C for 15 minutes, followed by inactivation at 65°C for 10 minutes. SPCs were then washed with WB three times, and to verify the occupancy of amplified genomes, SPCs were stained with 10x SYTO9 nucleic acid dye to observe amplification under a fluorescence microscope. SPCs were processed for DNA debranching and DNA end preparation as per the instructions in the manual.

### Primary template amplification

PTA was performed using the ResolveDNA Whole Genome Single-Cell Core Kit (Cat #100954, BioSkryb Genomics, USA) using the manufacturer’s protocol. The SPCs were treated with the lysis mix and mixed at room temperature at 1400 rpm for 15 min. Reaction mix was added subsequently, followed by a brief mix at 1400 rpm for 1 min. The mixture was further aliquoted into PCR tubes and incubated in a thermal cycler at 30°C for 2.5 hrs, followed by heat inactivation at 65 °C for 5 min and held at 4 °C. Further, SPCs were processed for DNA end preparation as per the instructions in the Atrandi manual.

### Cell Barcoding

Four-step combinatorial split-and-pool barcoding was performed to label each SPC with a unique variant. The barcodes were added by ligation following the Atrandi barcoding user guide, yielding up to 24^4^ barcode combinations.

### Library preparation for sequencing

The barcoded SPCs were further treated with the Release reagent (Atrandi Biosciences), and the released DNA was purified using 0.8X AMPure XP paramagnetic beads (Cat #A63881, Beckman Coulter). The purified DNA was used for the final library preparation using NEBNext Ultra II FS DNA Library Prep Kit for Illumina (Cat #E7805S, NEB) and custom PCR indexing primers. The library was cleaned using double-sided AMPure beads purification, and library concentration was measured by Qubit using a dsDNA HS kit (Cat #Q32854, Thermo Fisher Scientific), followed by checking the library QC on an Agilent Bioanalyzer. The libraries were further diluted as per the manual specifications and sequenced on an Illumina NovaSeq X platform.

### Analysis of scMetaG data from saliva

After debarcoding and trimming (i.e., using Bascet/Zorn as described above), reads were aligned (using the Bascet Rust translation of BWA-MEM2, default settings) to a concatenated reference genome consisting of human (T2T-CHM13v2.0^68^) and *Xanthomonas campestris* B-1459 (per the addition of xanthan gum in the scMetaG protocol; NCBI Assembly accession GCF_001372255.1). Reads that mapped to either genome were conservatively removed for raw data deposition and further analysis. To analyze the fraction of reads mapping to either reference genome, reads were considered to be mapping if the BAM unmapped flag (0x4) was clear and at least 90% of its read bases were mapped (CIGAR M).

Bascet/Zorn’s *k*-mer database workflow was used to obtain an overview of the sample using Kraken2 (*translated)* and the standard-8 database. Low-count filtering was performed, resulting in 10k cells used in subsequent analyses. SKESA was used to assemble SAGs *de novo* (*translated*) using parameters ‘--min_contig 50 --min_count 1--fraction 0.01’. QUAST-compliant QC parameters for SAGs were computed using Bascet.

### Bulk shotgun metagenomics of the saliva sample

250 μl of the processed saliva sample was used for the isolation of metagenomic DNA using the QIamp DNA Microbiome Kit (Cat #51704, Qiagen), with library preparation carried out using the Illumina DNA prep kit (Cat #20060059, Illumina). The library was sequenced using the paired-end run configuration of 2x150bp. Resulting reads were trimmed with fastp v0.23.4^49^ (default settings). The number of reads per species was estimated using Kraken2 (2.1.2) and the same database as used for the *k*-mer database workflow (which wraps a Rust translation of Kraken2). No reads were identified as “*Tannerella* sp. oral taxon 808” (NCBI Taxonomy ID 712711; **Supplemental Fig S2**).

### Host genome removal

Host genome removal was performed for saliva bulk and single-cell data by alignment using BWA-MEM2 (separate for bulk, v2.2.1, and *translated* for single-cell). Reads were considered “mapped” if (1) the SAM/BAM unmapped flag was not set, and (2) their CIGAR had enough M bases to cover 90% of the read sequence length. Mapped reads were removed. For the human mapped read percentage analysis, the mapped read percentage was simply based on the number of reads aligning to any human chromosome.

### Analysis of *Tannerella* sp. oral taxon 808 (TOT808)

Human saliva scMetaG reads (see sections “Saliva sample collection and processing” and “Analysis of scMetaG data from saliva” above) were aligned (Bascet BWA-MEM2) to the SAG of TOT808 (NCBI Assembly accession GCA_002890575.1).^28^ The Bascet *chromcount* command was used to produce a count table (chromosome vs cell) and the reads from cells with >10k reads were co-assembled using SPAdes v4.3.0^69^ (parameters: ‘--sc’).

The resulting co-assembled SAG was compared to all publicly available TOT808 genomes, metagenome-assembled genomes (MAGs), and SAGs. Briefly, GenBank versions of all genomes/MAGs/SAGs assigned to NCBI’s “*Tannerella* sp. oral taxon 808” taxon (NCBI Taxonomy ID 712711) were downloaded from NCBI’s Genome database (*n* = 3; https://www.ncbi.nlm.nih.gov/datasets/genome/, accessed 7 June 2026, **Supplemental Table S1**).^70^ The Genome Taxonomy Database (GTDB; v232, https://gtdb.ecogenomic.org/, accessed 7 June 2026) was subsequently used to identify additional TOT808 genomes/MAGs/SAGs, which had been submitted to NCBI under a different NCBI Taxonomy ID; GenBank versions of all genomes/MAGs/SAGs assigned to “*Tannerella* sp002890585” (GTDB’s alias for TOT808) were then downloaded from NCBI’s Genome database (*n* = 4 additional genomes, **Supplemental Table S1**).^71^ Finally, the mOTUs database (https://motus-db.org/, accessed 7 June 2026) was used to download TOT808 genomes/MAGs/SAGs (i.e., listed in the mOTUs database as species “*Tannerella* sp002890585”), which were not included in NCBI (*n* = 27 additional MAGs; **Supplemental Table S1**).^72^ Altogether, this search resulted in 34 total TOT808 genomes/MAGs/SAGs, which were used in subsequent steps (**Supplemental Table S1**).

All 35 TOT808 genomes/MAGs/SAGs (i.e., the 34 publicly available genomes/MAGs/SAGs, plus the co-assembled SAG generated here) were confirmed to belong to *Tannerella* sp. oral taxon 808/*Tannerella* sp002890585 using GTDB-Tk v2.6.1 (using ‘gtdbtk classify_wf’, version r226 of the GTDB database, and 28 CPUs; **Supplemental Table S1**).^73^ Genome/MAG/SAG quality was evaluated using CheckM v1.2.2 (’checkm lineage_wf -t 28 -f checkm_out.txt --tab_tablè)^74^ and QUAST v5.2.0 (default settings plus ‘--min-contig 1’; **Supplemental Table S1**).^75^ Average nucleotide identity (ANI) values were calculated between all TOT808 genomes/MAGs/SAGs using skani v0.1.4^76^ (’skani dist -t 1’; if either query or reference had QUAST N50 <3Kb or <10Kb, the ‘--slow’ and ‘--medium’ presets were additionally used, respectively, per the skani manual; **Supplemental Fig S3, Supplemental Table S2**).^76^

The co-assembled TOT808 SAG generated in this study, plus all high-quality TOT808 genomes/MAGs/SAGs (i.e., CheckM Completeness ≥80% and CheckM Contamination ≤10%; *n* = 9 total TOT808 genomes/MAGs/SAGs) underwent annotation using Prokka v1.14.6 (’prokka --kingdom Bacteria --cpus 4 --addgenes --centre X --compliant’).^77^ The resulting GFF files were supplied as input to Panaroo v1.3.4 (’panaroo --clean-mode strict -a core --aligner mafft --core_threshold 0.95 -t 28 --remove-invalid-genes -f 0.5’).^78^ The filtered core gene alignment produced by Panaroo (’core_gene_alignment_filtered.aln’), which consisted of 103 core genes detected in all 9 TOT808 genomes/MAGs/SAGs, was supplied as input to IQ-TREE v2.2.5^79^ and used to construct a maximum likelihood phylogeny, using one thousand ultrafast bootstrap replicates (’-B 1000’)^80^ and the best-fitting nucleotide substitution model selected using ModelFinder (’-m MFP’; model ‘TIM2+F+R2’).^81–83^ The resulting maximum likelihood phylogeny was rooted at the midpoint (using the ‘midpoint.root’ function from the phytools v2.5-2 R package)^84^ and visualized in R v4.6.1, using the ggtree v4.2.0 package (**Fig 4f**).^85^

The aforementioned genome annotation, core gene alignment, and maximum likelihood phylogeny construction/visualization process was repeated, with the addition of an outgroup genome (i.e., *Tannerella serpentiformis* strain W11667; NCBI Assembly accession GCF_001717525.2), resulting in the following deviations from the process described above: (i) the core gene alignment produced by Panaroo consisted of 22 genes; (ii) the best-fitting nucleotide substitution model selected by IQ-TREE/ModelFinder was the ‘TIM3+F+I+R2’ model; (iii) the phylogeny was rooted at the outgroup, using the ‘root’ function from the ape v5.8-1 R package (with ‘resolve.root = T’; **Supplemental Fig S4**).^86^

### Code availability

Bascet and Zorn are both open-source and freely available via GitHub under the MIT license (https://github.com/henriksson-lab/bascet and https://github.com/henriksson-lab/zorn, respectively). Code specific for this study has been deposited at GitHub (https://github.com/henriksson-lab/scwgs2026). scMetaG and bulk metagenomic data for the saliva sample are available at Biostudies #S-BSST3416 (host removed). The mock community scMetaG data is available at Biostudies #S-BSST3417. The simulated mock community is available on Zenodo https://doi.org/10.5281/zenodo.15813069.

## Supplemental Figures

**Supplemental Figure S1.**
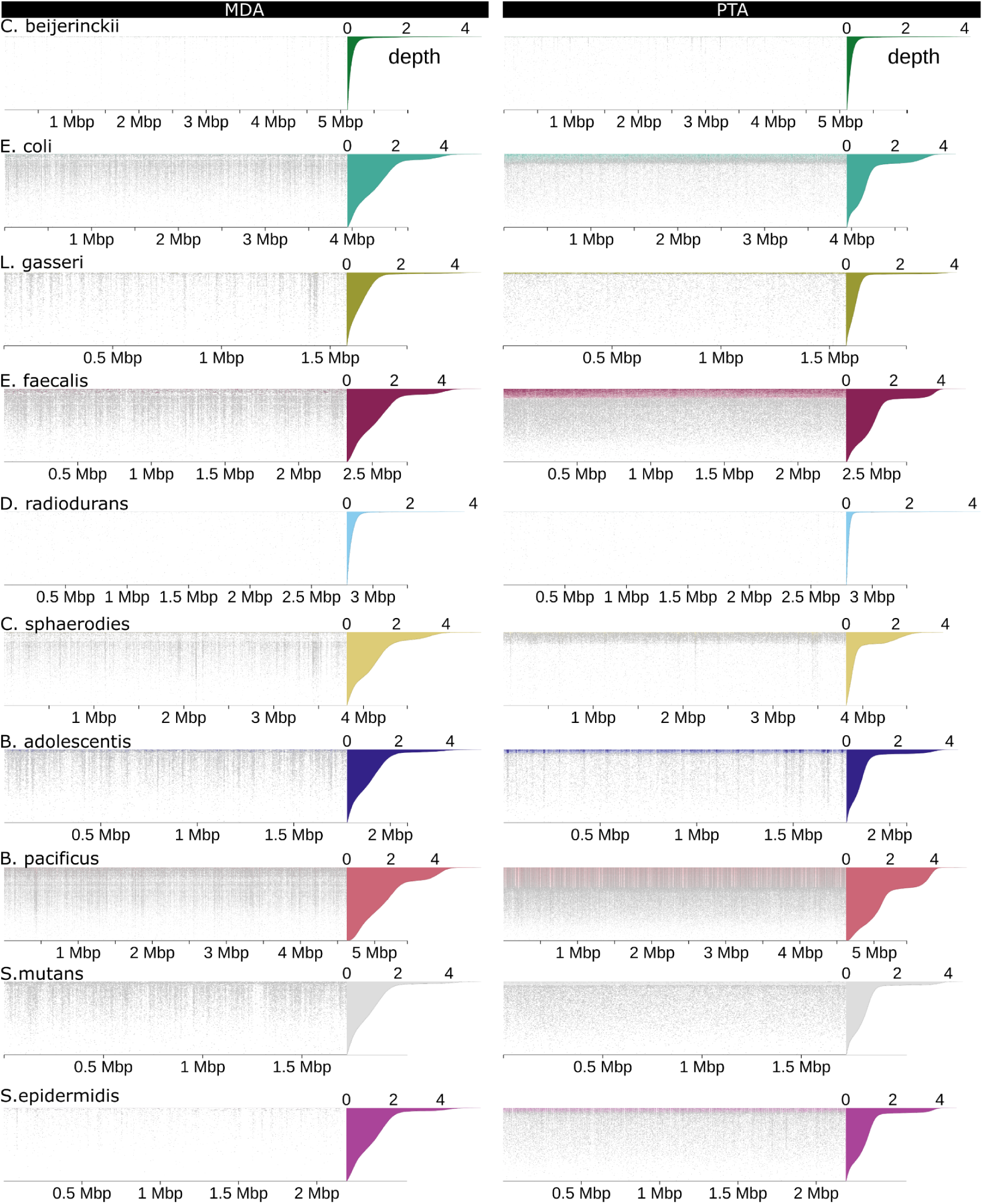
Pileups of all species in the mock community, for MDA and PTA. Each row is one cell. Only top-ranked cells for each species are shown. All cells for B. pacificus are shown in Fig 2e.

**Supplemental Figure S2:**
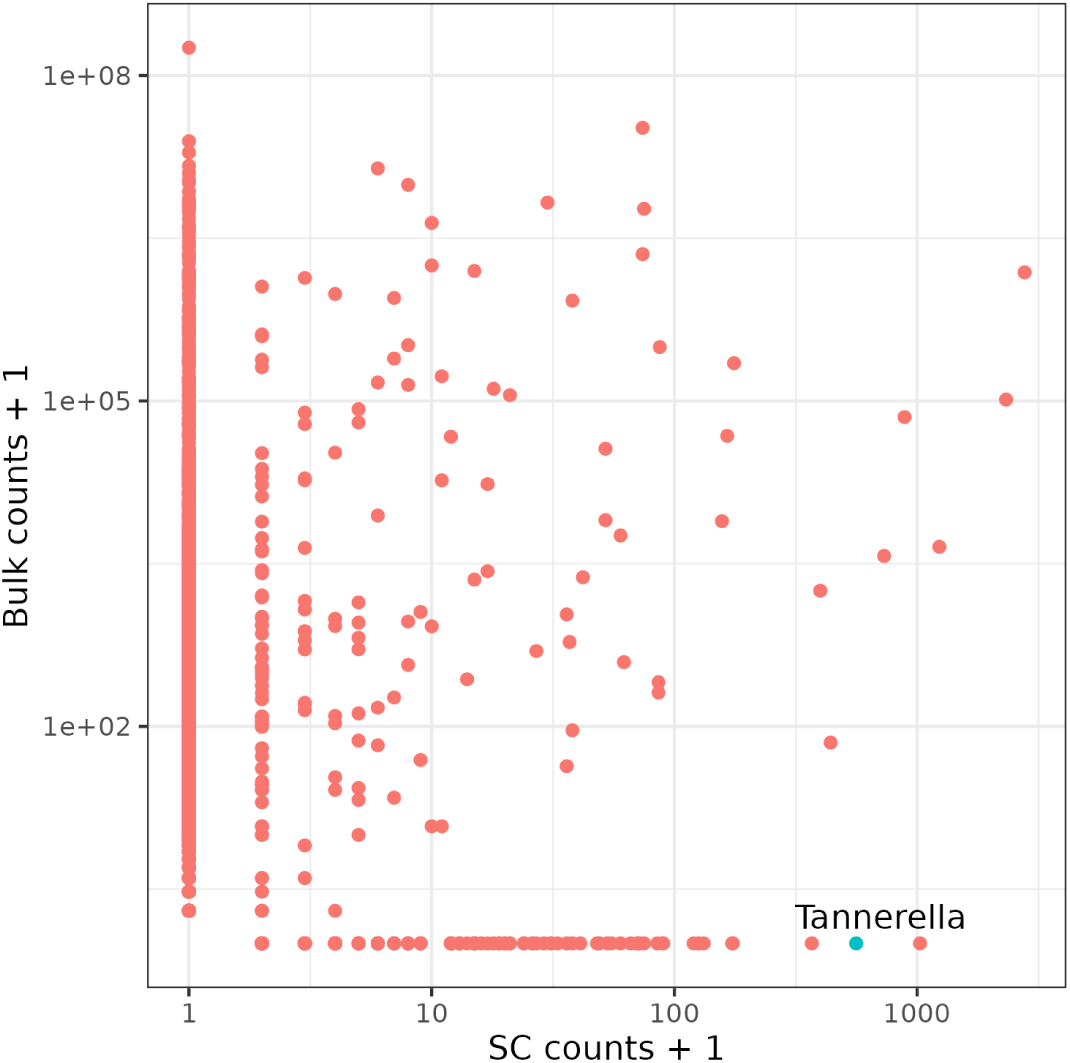
Comparison of read count abundance for Tannerella sp. oral taxon 808 (TOT808; blue dot with “Tannerella” text label), for saliva single-cell data (X-axis) vs bulk data (Y-axis) after host removal.

**Supplemental Figure S3.**
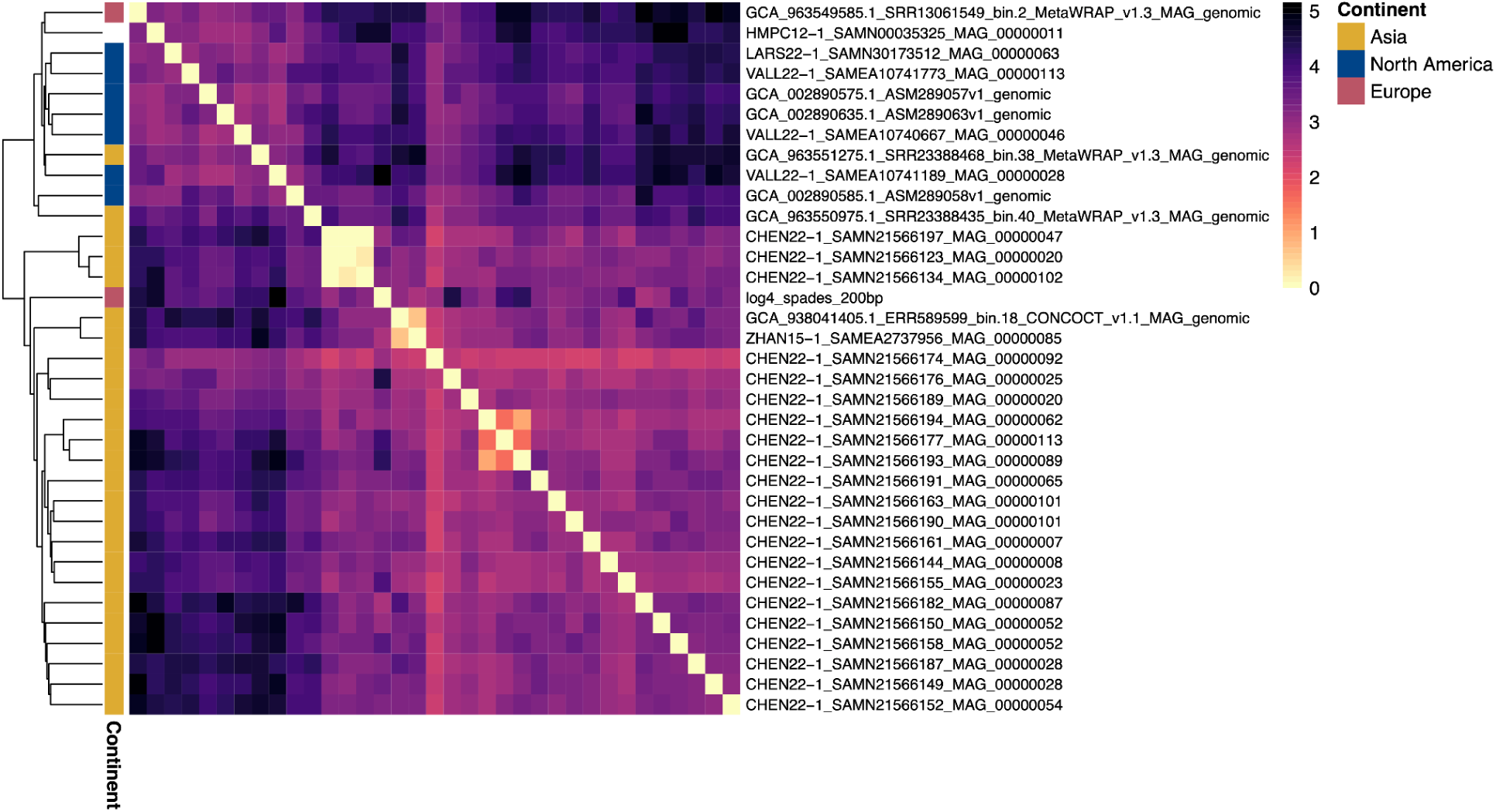
Heatmap of pairwise average nucleotide identity (ANI) dissimilarity values (100-ANI) calculated between all Tannerella sp. oral taxon 808/Tannerella sp002890585 genomes using skani.^76^ The color strip to the left of the heatmap shows the continent of origin for each genome. The dendrogram was produced via average linkage hierarchical clustering using ANI dissimilarities (100-ANI). The heatmap was constructed using the ‘pheatmap’ function from the pheatmap v1.0.13 R package.^87^

**Supplemental Figure S4.**
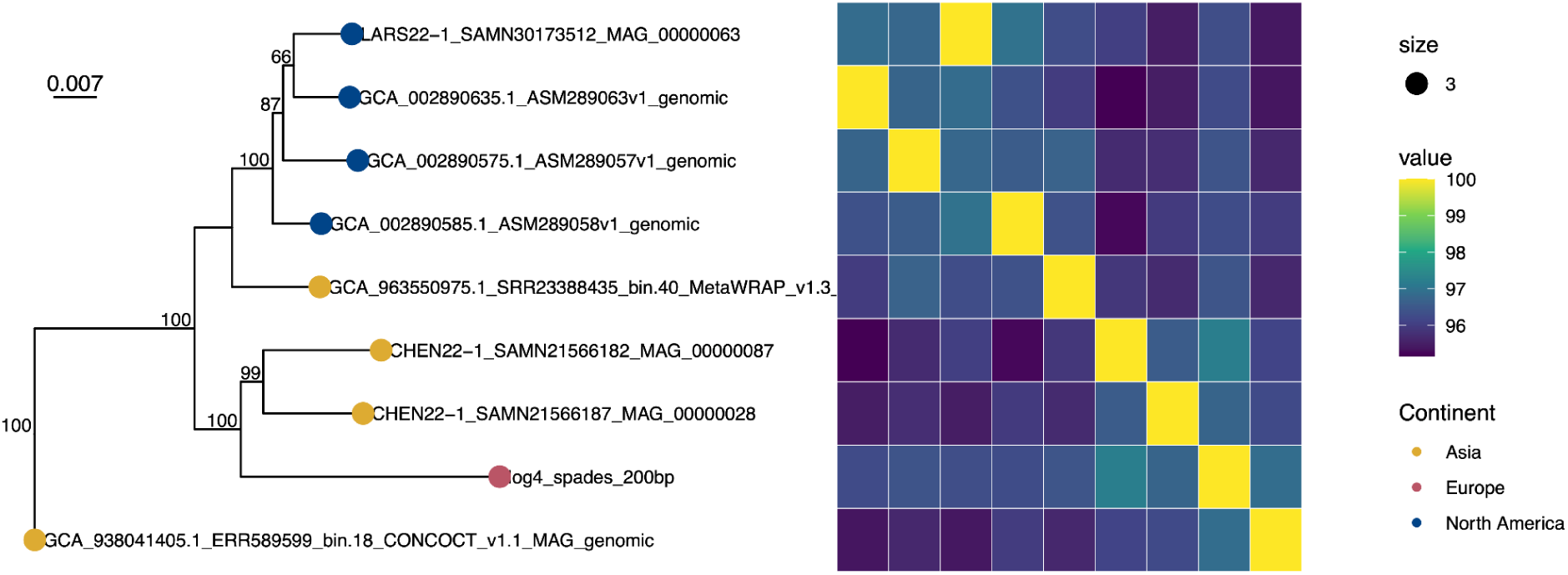
Maximum likelihood phylogeny constructed using an alignment of 22 core genes detected in all high-quality Tannerella sp. oral taxon 808 (TOT808) metagenome-assembled genomes (MAGs)/SAGs, plus a novel TOT808 SAG generated here (tip label log4_spades_200bp) and an outgroup (Tannerella serpentiformis strain W11667, NCBI Assembly accession GCF_001717525.2). The phylogeny is rooted at the outgroup (omitted for readability). Branch lengths are reported in substitutions/site. Branch labels denote IQ-TREE ultrafast bootstrap support percentages. Colored tip shapes (circles) denote the continent from which each sample was reportedly collected. Average nucleotide identity (ANI) values calculated between each MAG/SAG are displayed in the heatmap to the right of the phylogeny.

**Supplemental Figure S5.**
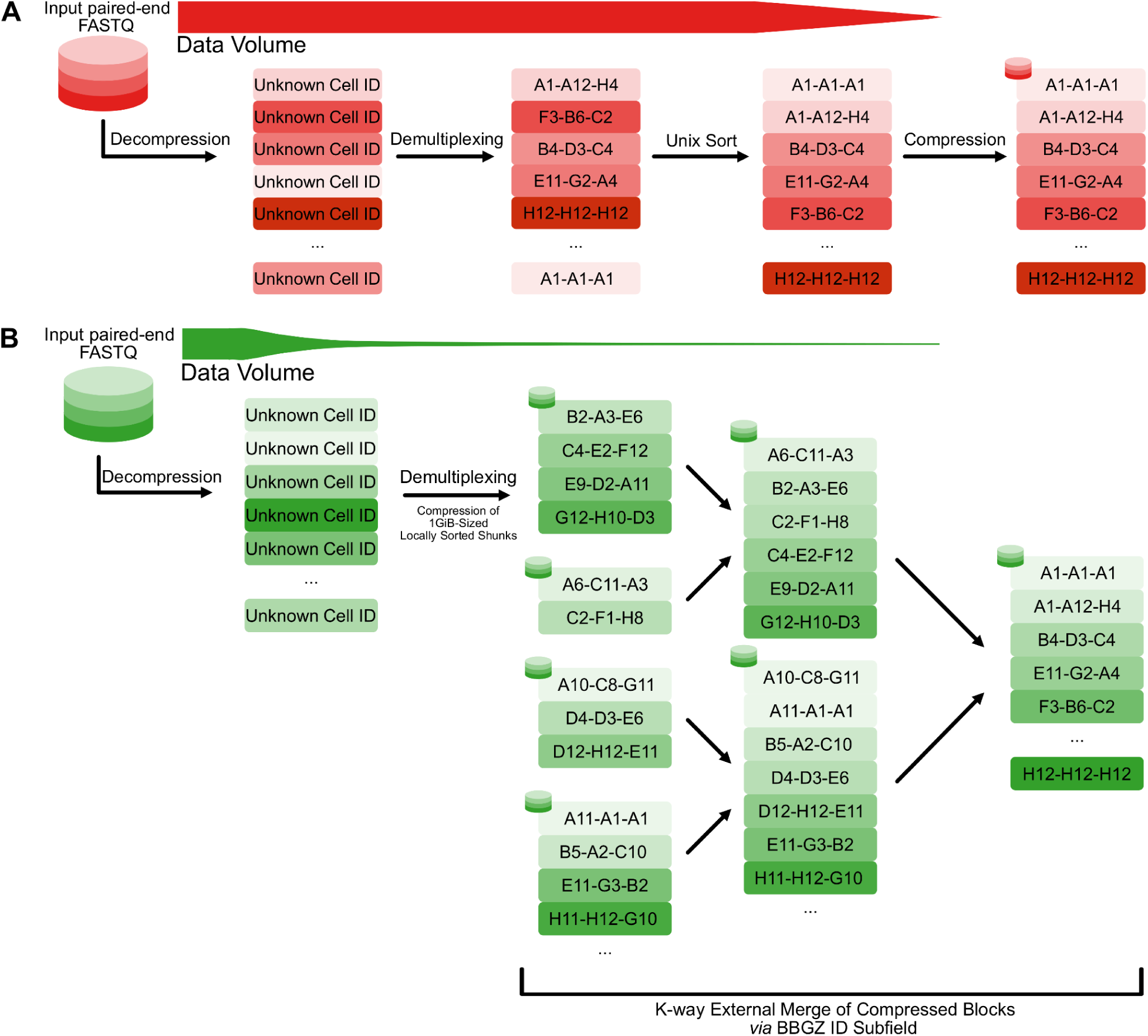
Comparison of conventional and BBGZ-based single-cell data preprocessing workflows. **(a)** Conventional workflow: paired-end FASTQ files are fully decompressed, demultiplexed by cell identity, sorted via Unix sort on the decompressed data, and recompressed. Data volume (red bar) remains high throughout, as all operations must act on decompressed data. **(b)** Bascet block gzip (BBGZ) workflow: reads are decompressed and demultiplexed into locally sorted chunks of approximately 1 GiB and then compressed. These chunks are then merged into a globally sorted output via K-way external merge using only the BBGZ ID subfield, without decompression. Data volume (green bar) is reduced as sorting and merging operate on compressed blocks. Compressed data is indicated by a database symbol.

**Supplemental Figure S6.**
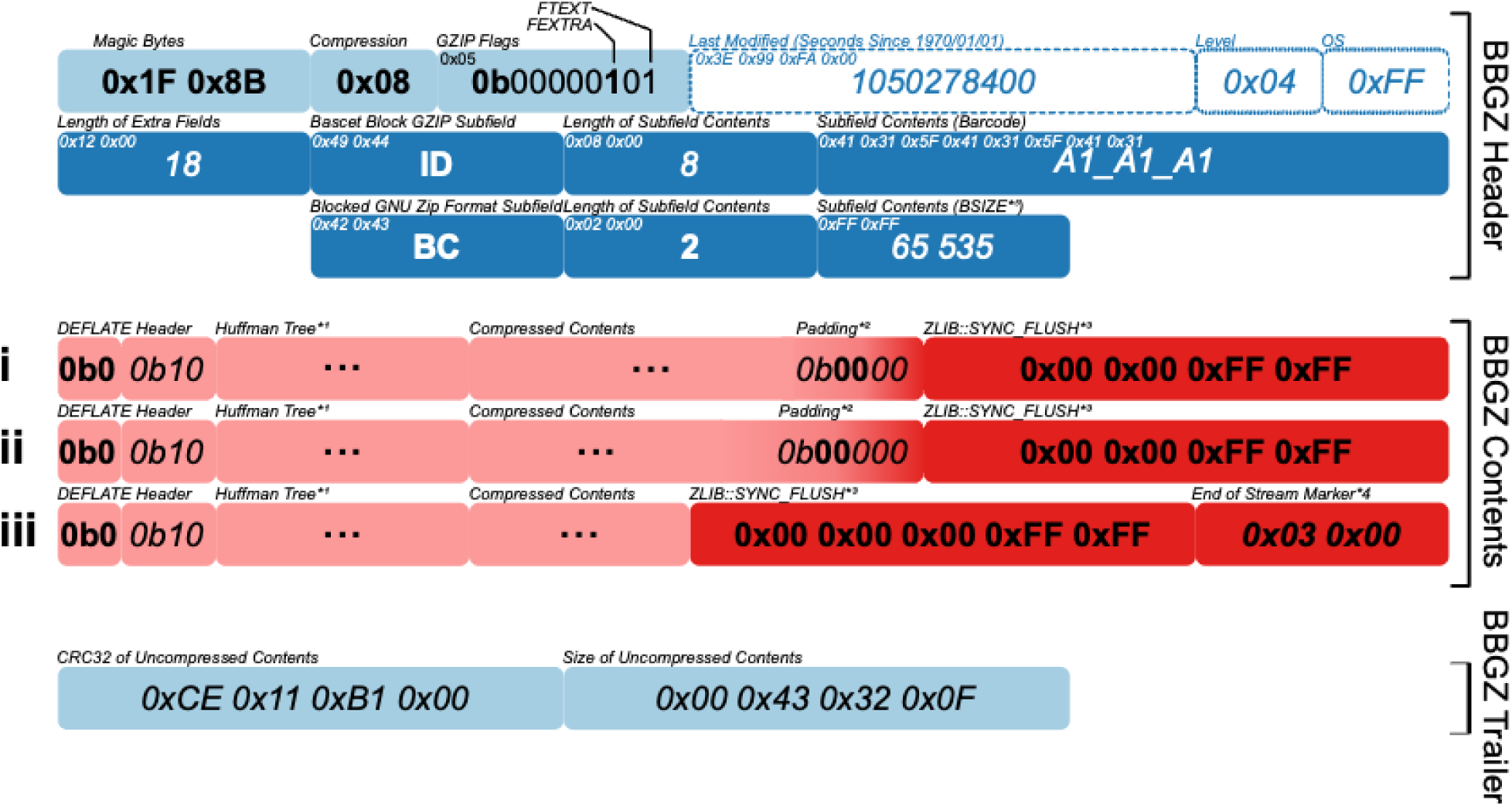
Structure of a BBGZ block. The GZIP header contains two subfields: an ‘ID’ subfield encoding the cell barcode (here “A1_A1_A1”) and the standard BGZF ‘BC’ subfield required. The compressed contents consist of multiple DEFLATE blocks (i–iii), each terminated by a ZLIB::SYNC_FLUSH byte-alignment marker that may be of varying bit length (i-ii) or omitted in the case of a naturally byte-aligned bit stream (iii). The final block (iii) is followed by an end-of-stream marker. The block concludes with a GZIP trailer containing the CRC32 checksum and uncompressed size to verify data integrity. Bold values are constants, italic values are examples, and hollow fields are mandatory GZIP fields with no functional role in the BBGZ format. **Remarks:** *1: Huffman tree representation per DEFLATE specification. *2: To facilitate parallel (de)compression of DEFLATE streams, a DEFLATE block must be byte-aligned. Padding bits align DEFLATE streams to the nearest byte-boundary. *3: A literal empty DEFLATE block written via ZLIB::SYNC_FLUSH to align the DEFLATE bit stream to the nearest byte boundary. The literal DEFLATE block flag may be contained in the most significant bits of the padding bits or within the sync flush marker, depending on the byte alignment of the underlying DEFLATE stream. *4: The end of stream marker to signal “End of Data” to the decompressor.

