## Supplemental Note for "Scalable single-cell metagenomic analysis with Bascet and Zorn"

#### Small files cause performance problems

Common software for scRNAseq analysis (e.g., 10x Genomics CellRanger, Parse Biosciences split-pipe, STAR solo[1], zUMI[2], etc.) bundle cells together in single large files to gain efficiency. The typical raw data storage format for scRNAseq or scATACseq is 3 FASTQs, including read1 (R1), read2 (R2) and index 1 (I1). This is obtained by running Illumina bcl2fastq (image-to-read demultiplexer) with NNNNNN set as the library index, and providing the flag --create-fastq-for-index-reads. Cell barcodes are then identified (which are commonly in a read rather than the index), and the reads fed to an aligner. After alignment, cell barcodes are kept as special tags in the resulting BAM file (for CellRanger, the output format is specified at https://www.10xgenomics.com/support/software/cell-ranger/7.2/analysis/outputs/cr-outputs-bam) . After sorting the BAM file by coordinate, features can be counted for all cells in parallel (**Fig 1bc** of the main manuscript).

The CellRanger approach works well for scRNAseq data, but does not support the needs of *de novo* assembly, where the gathering of reads belonging to a single cell must be fast. It also relies on the presence of a reference genome, which is typically not the case for metagenomics (**Fig 1bc** of the main manuscript). Storing all reads in individual FASTQ files would solve the gathering problem, but causes a great number of problems due to the way operating systems, file systems, network protocols, and software are designed (**Sup Note Fig 1**). This section lists a number of specific issues and also benchmarks a few cases.

#### Transfer speed is reduced due to handshakes

The transfer of a file using a protocol such as SCP, FTP or NFS is shown in **Sup Note Fig 1e**. The operation can be thought of as happening in two steps: (1) The user on the client computer requests a file from the server, and (2) the server sends the content of the file. The time this operation takes depends on the network connection, but the performance of the connection can be described by two parameters. The time it takes for a message to reach from a-to-b is called *latency*. For internet connection, the latency can be checked by the command *ping* (e.g., the time from home office to our cluster, HPC2N, is 10ms). The other parameter is *bandwidth*, which for a good internal network is 1 gb/s, but may be much higher for computing clusters using infiniband connections or advanced solutions. Latency need not be correlated to bandwidth as the time from a-to-b is limited by physics (e.g., the speed of light, or electrons in copper). However, a large number of parallel channels can be used to transfer data back once connection has been made, achieving high bandwidth despite possibly high latency.

Latency can be a serious limitation for transfer speed. As an extreme example, specialized network protocols have been designed to send data to other planets, where latency can be in the range of minutes (delay-tolerant networking, CCSDS being one protocol for the purpose https://ccsds.org/wp-content/uploads/gravity_forms/5-448e85c647331d9cbaf66c096458bdd5/20

25/06/734x20o1.pdf). If files are requested one at a time, and the files are small, latency will dominate overall transfer time (**Sup Note Fig 1e**). The solution for CCSDS is to bundle a large number of requests into one message, such that correspondingly, a large number of files can be sent back in one round. Similarly, copying many small files using SCP is much faster if the files are first tar-ed (concatenated) on server side such that only a single file needs to be requested.

While network communication has been used as an example here, computers are all networks at multiple levels; e.g., the link between CPU and main memory is a network, and similarly, the link between CPU and internal computer hard drives. Small files thus induce communication overhead whenever many files are requested, but only one at a time. Since this is how current operating systems are designed,[3,4] the only way to circumvent multiple requests is through concatenating the files into single larger files.

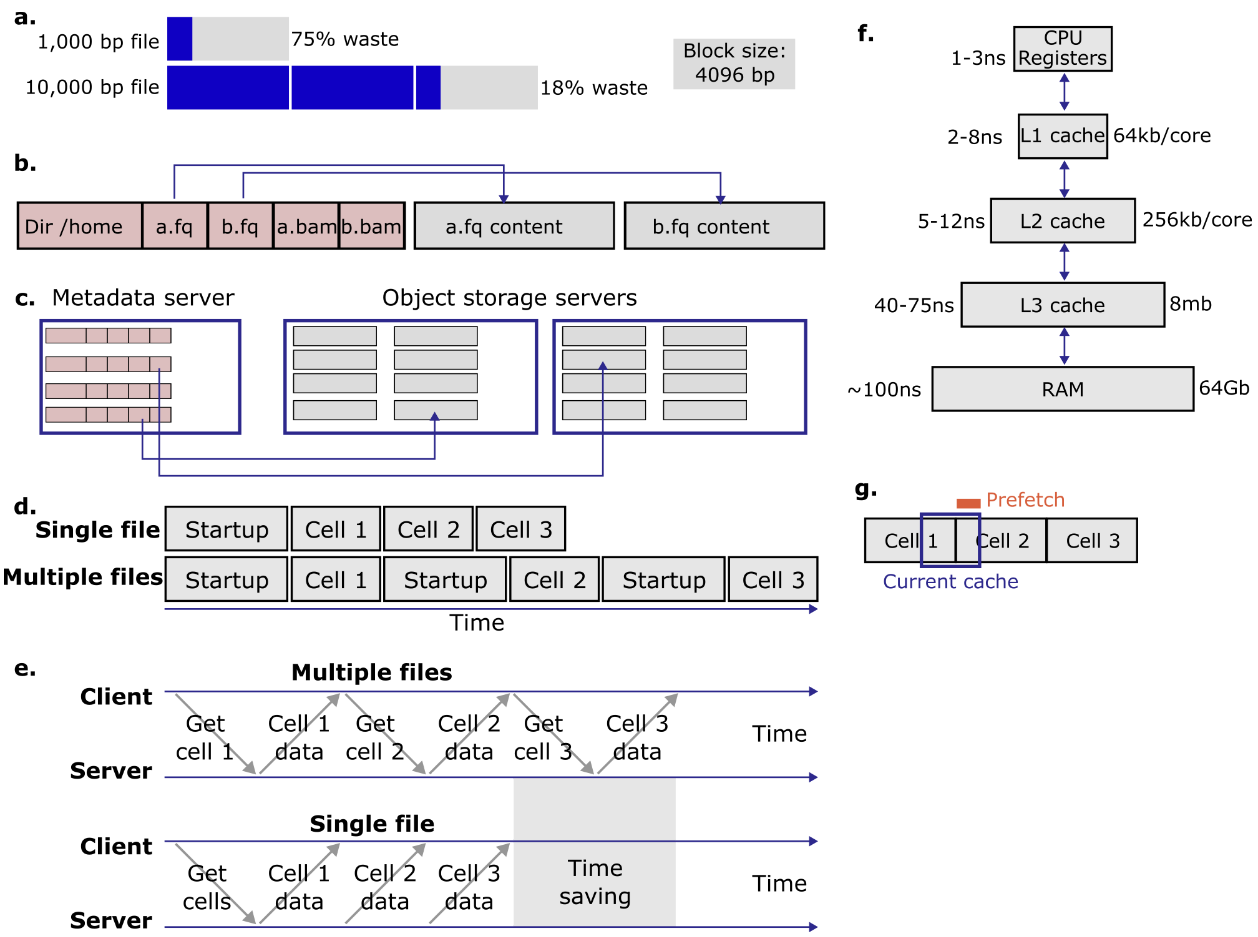

**Supplemental Note Figure 1: Small files cause performance problems. (a)** Block-based allocation of files causes small files to waste memory. **(b)** File systems store file metadata separately from file content. **(c)** On network file systems, metadata is commonly stored separately from file content, turning frequent metadata lookup into a bottleneck. **(d)** Repeated software startup time results in overhead for processing many files. **(e)** Network file transfer is bottlenecked by latency time if many files are fetched. **(f).** Memory access time is a major limiting factor for modern computers, requiring data to be laid out in a cache-optimal manner.

Times and sizes are approximates for the Intel skylake family. **(g)** Data prefetch systems, and cache memories, benefit from data used being laid out consecutively in memory (ram and disk).

#### Small files are difficult to compress efficiently

It is common to work with compressed files, where especially FASTQ files benefit greatly. Compression works by increasing the entropy of the stored content, or in other words, reducing repetition. As an example, the binary ASCII representation of A is b01000001, and for T it is b01010100. The sequence AATT is thus b01000001010000010101010001010100. However, if we instead create a specialized dictionary, such that A=b0, and B=b1, then the AATT can simply be represented as b0011. This is a reduction from 32 bits to 4 bits (8x compression).

For simplicity, we will only consider the example of dictionary-based compression schemes, as previously illustrated. In a simple version, the string AATT would be stored in two parts: {dictionary}{compressed data}. If the dictionary is *small* and can be used for a *large* amount of data, the result is that the dictionary and compressed representation will occupy less space in total than the original text.

The example illustrates a general trend that larger amounts of data can be compressed better than smaller amounts of data, as a dictionary (or equivalent) must be built and stored as well. A big file with reads from 2 cells is compared to two cells in separate files in **Sup Note Fig 2**. Although the name of the cell a read belongs to has to be repeated in the big file, this content is highly repetitive and easy to compress. Once a compressor has figured out a suitable dictionary for the DNA, the dictionary can be used for the entire file. For the second scenario, the dictionary has to be rebuilt for each file, preventing it from warming up on small files. Thus, small files benefit less from compression.

Tricks exist to mitigate the problem of small files, e.g., dictionaries can be built/"trained" from large amounts of data, even if only used for a smaller file of the same type; or the dictionary can be stored in a separate common file. However, these approaches require developer intervention and generally prevent use of off-the-shelf compressed file formats.

| | |
|---|---|
| **BIG FILE**<br>>cell1<br>ACAGTACGATCGATCGTACGATCGATCG<br>>cell1<br>ACGACGAGCATGATCGATCGATGCATCG<br>>cell2<br>CGATCGATCAGATCGATCGATCAGTACG<br>>cell2<br>CAGTCAGTAGATCGATCGATCGTAGCTA<br><br>Gzip ⇒ **91 bytes** | **FILE 1**<br>><br>ACAGTACGATCGATCGTACGATCGATCG<br>><br>ACGACGAGCATGATCGATCGATGCATCG<br><br>**FILE 2**<br>><br>CGATCGATCAGATCGATCGATCAGTACG<br>><br>CAGTCAGTAGATCGATCGATCGTAGCTA<br><br>Gzip ⇒ 61+60 = **121 bytes** |

**Supplemental Note Figure 2: Small files benefit less from compression.** A big file with reads from 2 cells (BIG FILE, left), versus two cells in separate files (FILE 1 and FILE 2, right). Gzip (Apple gzip 479) performs better on the big file despite having more text in total. This is a general trend for any choice of compression of software.

#### Space is wasted due to file system organization

File systems are typically organized around blocks, from 4kb to 64kb. A file occupies an even number of blocks and starts at the beginning of a block. This reduces the size of addresses to the starting position of files by at least 12 bits/file and reduces index size. However, hard drive physical interfaces are also addressed in terms of logical blocks (512 bytes), and attempts to work at finer resolution typically result in reduced performance (i.e, the full block has to be read in either case). For this reason, small files have always been a problem for file systems. The storage of a file in a classical file system such as FAT32 (Windows) or Ext3/4 (Linux) is shown in **Sup Note Fig 1a**. As the allocated file size is rounded up to even block sizes, small files incur larger losses. A FASTQ file with a single 150bp read (~350 bytes), for example, will waste 91% of the allocated space ($1 - 350 / 2^{12} = 0.915$). Exceptions to the allocation rule exist; some newer file systems can store small files directly in sub-blocks or directly in the metadata table for the file. ReiserFS was among the first to do it, but NTFS (files that are < 900 bytes) and Btrfs (files < 2kb, configurable) are also capable. Designing pipelines that can only perform well on the right choice of file system is poor design, especially as users of computing facilities cannot influence this decision. Storage of file content in the file metadata area is also poorly compatible with networked file systems.

#### Network file systems are especially sensitive to small files

File systems can typically be thought of as separated in two areas, shown in **Sup Note Fig 1b**: (1) metadata and (2) file content. The metadata holds information about directories and their contents, including dates and permissions. The metadata block then points to the area on the disk where the file is actually stored. As metadata has to be stored for files, they also occupy a fixed amount of additional data in the metadata partition, independent of their size.

Metadata storage is a particular bottleneck in computing facilities. Lustre (https://www.lustre.org/) is a common file system that illustrates the problem. In Lustre, metadata is kept on separate servers from those serving the file content (**Sup Note Fig 1c**). This design is motivated by the rather different access patterns of metadata *vs* file content, where metadata management is a rather complex, fine-grained process, while file content is commonly just streamed from start-to-end.

As metadata is stored on a small number of metadata servers, they can become a weak link when this architecture is put under load. It is, e.g., common to disable coloring of files when using the ls command to list files in Unix (--color). Coloring a file, depending on if it is a directory or not, requires reading this additional metadata, which adds extra load. If a user requests information about a large number of files, it may overload the metadata server to the point where other users are put in queue to obtain the information they need. A single user may thus bring down a whole computing cluster if trying to access a large number of small files.

#### Large numbers of jobs may crash SLURM

SLURM (https://slurm.schedmd.com/) is one of the most common job managers, responsible for executing computational tasks on compute clusters. To operate it, a user writes and submits a BASH shell script to drive the computation, along with requests on the amount of memory, CPU cores, and other resources. SLURM then performs an optimization task aimed to maximize the resource utilization, i.e., running as many jobs as possible on given computers that can fit the total amount of memory and CPU cores requested. This optimization task is computationally intensive, and the SLURM server can thus be overloaded. In such a scenario, jobs can no longer be submitted, and the entire HPC cluster may have to be restarted, affecting all users.

SLURM is able to execute job arrays, i.e., a series of highly similar jobs. A typical use case is to perform *de novo* assembly of a number of genomes, with 123 genomes resulting in a job array of size 123. It is easy to create a large number of jobs using arrays, and thus overload SLURM; it is thus frequently configured to disallow large arrays. As an example, on the Swedish national computing cluster Dardel, the cap is at 1,001 jobs ("scontrol show config" ⇒ MaxArraySize). This makes it difficult to naively process more than 1,001 single-cells if stored in individual files.

A work-around is to submit *mini batches*, i.e., a job that may in turn process some 1000 files. In this manner, 1,001,000 single cells could be processed in individual files. However, this requires the user to effectively make their own work manager, in turn wrapped in SLURM (which takes time to implement and adds a level of complexity that may cause errors), or use additional specialized software for this task (e.g., Nextflow; https://www.nextflow.io/). To conclude, processing many small files is inefficient and difficult with SLURM.

#### Program initialization overhead is excessive for small files

Bioinformatics tools typically run in 3 phases: initialization, data processing, writing of results. If a tool requires reading of a database as part of initialization, this can add a large amount of startup time overhead (**Sup Note Fig 1d**). An example is the tool Kraken2, which first reads a database. Once loaded, reads can be processed, and results written at high speed.

To ensure high performance, Bascet sends the reads of all cells to Kraken2 in a single batch, requiring only a single loading of the database. Based on the runtime on our computer (Intel Xeon Gold 6138 2.00GHz, 192GB RAM; version 2.1.2 with --thread 10), we can estimate the runtimes of Bascet Kraken2 vs naive Kraken2 (i.e., running Kraken2 on one genome per input file; **Sup Note Fig 3**). We compare performance for the small 7.5 GB standard-8 index (obtained from https://benlangmead.github.io/aws-indexes/k2). The naive approach is 887x slower, as it has to keep loading the database, whereas Bascet keeps the database in memory across all cells (**Sup Note Fig 3**). The speed benefit increases for larger databases and startup overhead.

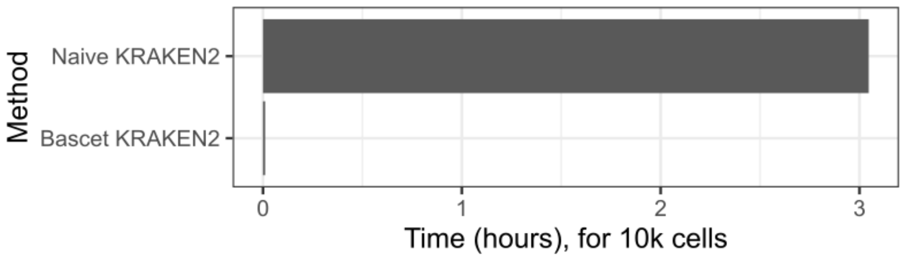

**Supplemental Note Figure 3: Performance of Kraken2**, run on a naive file-by-file (i.e., genome-by-genome) basis (Naive Kraken2, top bar), compared to Bascet-Kraken2 (bottom bar). Bars denote the time required to process 10k cells using the Standard-8 database.

Another example of startup overhead is when an aligner is used. Assuming BWA-MEM2[5] is used to align reads to the human genome (to remove host DNA), we estimate BWA-MEM2 startup cost to be 11.1 s, compared to the alignment speed of 0.61 s/cell. This means 94.7% of the time is spent starting the software when run cell-by-cell (**Sup Note Fig 4**).

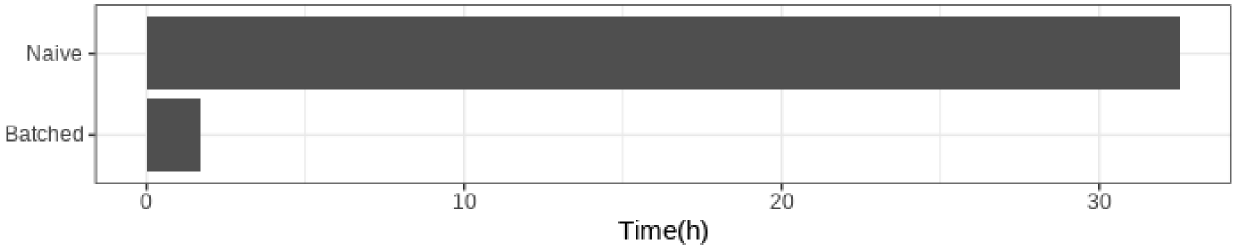

**Supplemental Note Figure 4: BWA-MEM2 alignment time** for 10k cells (800M reads). **Naive**: Cells processed one at a time. **Batched**: All cells aligned in one sweep, as done by Bascet.

A way to reduce the program overhead is to effectively never shut it down between jobs. For example, the aligner STAR has an option (--genomeLoad) to share the genomic index between processes, and to retain it in memory after the program has shut down. This option has to be specifically coded and may backfire if the user forgets to unload the genome. While it may be useful in certain scenarios, it breaks the idea that a program ends when it stops executing, adding complexity and making the workflow fragile. Furthermore, not all software has such an option. It is thus overall a bad design, only needed to overcome poorly managed input data.

#### BASH is not designed to handle large numbers of files

BASH is the most common tool to interact with the file system and run software on Linux. If more than 50,000 files are stored in a directory and you wish to delete them, you will likely encounter the following error message:

```
b-an01 [~/directory_with_many_files]$ rm *
bash: /usr/bin/ls: Argument list too long
```

This error is because BASH will first expand * to a list of all files in the directory. For 50,000 files, each file name being 20 letters, this results in a command that is more than 1,050,000 letters.

BASH is not designed for long commands and may instead give an error. On POSIX-compliant Unix systems, the limit can be checked by the command *getconf ARG_MAX*. On OSX 15.5, the limit is 1,048,576; on Linux, the limit varies depending on the distribution and administrator setting. As a result, working with directories having many files is difficult, and the precise number is user-dependent in a hard-to-predict manner.

Similarly, the maximum content in environment variables is limited. Linux has had limitations between 128kb and 8mb, depending on version. This limitation is for all environment variables combined, making the limit for a single variable hard to predict.

Software can in general operate with directories having a large number of files. However, because many bioinformatics tools are designed to be run by the BASH command line, BASH limitations may in turn be imposed on other software, e.g., if they expect a list of input files as command line arguments, or in an environment variable. Working with software that expects data from individual cells to be stored in separate files can thus be challenging.

#### Small files induce cache misses and context switches

In a modern computer, the CPU and especially the transfer of data from RAM to the CPU has become a major bottleneck, as the speed increase of these components has stagnated. It is thus essential to address this bottleneck at an early stage of the design of new software.

Because accessing RAM from the CPU is slow, the CPU is rather accessing it through a series of faster cache memories (L1-L3; L4 is sometimes also present; **Sup Note Fig 1f**). A key idea is that memory access tends to be *local*, and thus, by loading an entire segment of memory from RAM into faster local memory, the access time for subsequent memory access can be reduced (see analog discussion about latency *vs* bandwidth in previous section). The difference in speed in L1 cache *vs* the RAM can be up to a factor 100x, making this likely the most important optimization for software performance. Many detailed considerations (and benchmarking) must be considered when optimizing cache memory access; however, the benefit of concatenating small files can be seen in **Sup Note Fig 1g**. When memory is read from RAM to L3 cache, L3 to L2, and so on, the hardware will optimistically *prefetch* subsequent memory. If the data for the next cell to be processed is located directly after the current cell, prefetching will result in required data being loaded immediately. Similarly, hard drives also have cache memories and perform prefetching, and the advantages thus ripple down to the source of the data.

Small files not only prevent prefetching but also interrupt the process at a deeper level. To process the next file, it must be opened, requiring a call to the operating system. Unlike a typical function call, this requires a *context switch*, in which the CPU is reconfigured for elevated access privileges but also for using a different memory address space. The context switch takes about 1.2-1.5us, or 1200-1500ns, to execute (https://eli.thegreenplace.net/2018/measuring-context-switching-and-memory-overheads-for-linux-threads/). The cost of context switches are thus 3 orders of magnitude higher than the cache misses we have aimed to address in the design of Bascet.

### Benchmarking of Bascet/Zorn RNA-seq performance

To our knowledge, there is no tool for scMetaG analysis that operates on concatenated cells stored in large files (e.g., as for scRNAseq data), and thus, there are no existing scMetaG analysis tools to benchmark Bascet/Zorn against. We instead benchmark Bascet/Zorn against CellRanger (10x Genomics) using *scRNAseq data*, as a subset of the Bascet/Zorn workflows are capable of also doing such analysis.

Briefly, we obtained 353 M human 10x Genomics scRNAseq reads[6] and used CellRanger 10.0 to align to its refdata-gex-GRCh38-2024-A. The same reference was used for Bascet. Both software packages were given 40 threads and 50GB RAM. The results are shown in **Sup Note Fig 5** and indicate that Bascet is 8x faster than CellRanger (wall time).

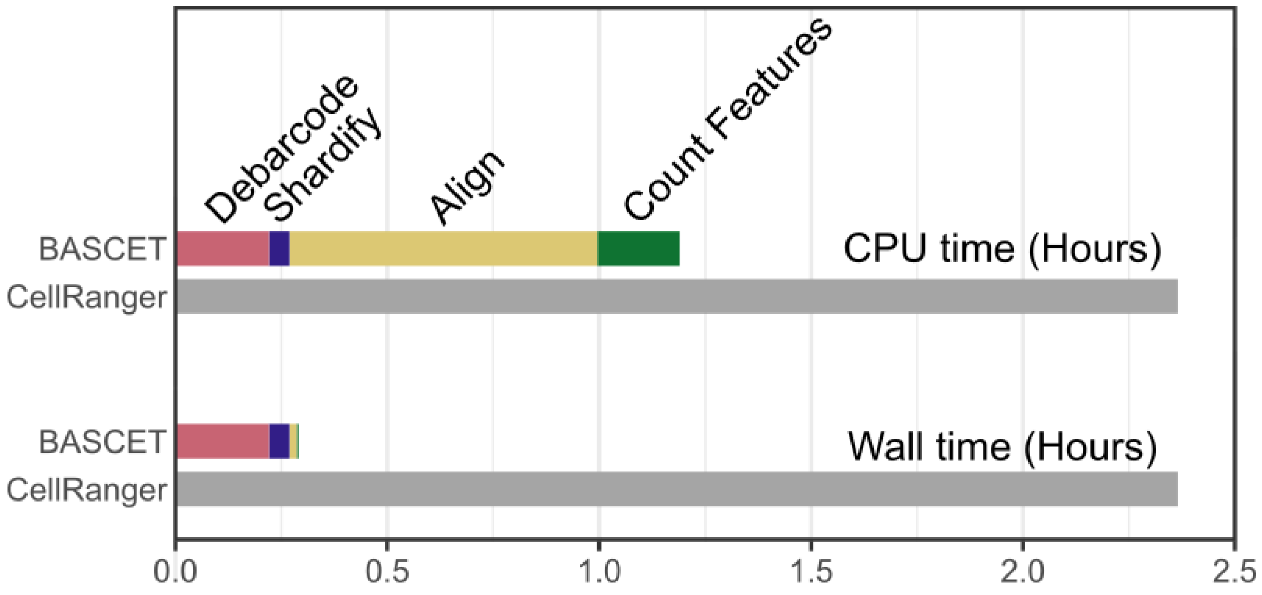

**Supplemental Note Figure 5: Performance of Bascet compared to the commercial 10x Genomics pipeline CellRanger**, for scRNAseq.

#### Bascet processing time on a mock community dataset

**Sup Note Fig 6a** shows Bascet's runtime on scMetaG data generated from an ATCC 10-strain mock community (generated using primary template-directed amplification [PTA]; see **Online Methods** for details). The work was performed at the Swedish facility HPC2N using nodes with AMD EPYC 7313 CPUs (only 20 cores were requested for the jobs). For simplicity, 100gb RAM was used for all steps (our facility has multiple generations of CPUs, and picking a high amount of RAM enforces use of the latest generation) but most steps will function with <32gb. Bascet performs faster with increasing RAM but can operate well on as little as 8 gb RAM. However, the first step, Debarcode, also performs initial rough sorting of reads, and by providing additional RAM, further sorting can be performed before storing reads on disk. Sorting speeds up the following Shardification, which uses the custom tags in the DEFLATE blocks to avoid uncompressing and recompressing reads (described in **Online Methods**).

We benchmarked Bascet on a dataset of **782,425,486 reads (800M, equivalent to 1 lane of a Novaseq X 1.5B).** The workflow includes raw read processing, host DNA filtering, creation of four types of UMAPs (alignment-based, informative k-mers, countsketch, and Kraken2-based) and assembly of single-amplified genomes (SAGs). 20 shards were produced, enabling up to 20 nodes to process the data in parallel (wall time is computed based on this assumption). Debarcoding is, however, typically limited by the number of input FASTQ files (commonly one pair). Shardification is also limited to one node, but most of the work (sorting) is done already during debarcoding. Thus, for larger jobs, with more input FASTQ files, shardification is not much of a bottleneck (**Sup Note Fig 6**).

Processing time on a single compute node (CPU time) is about 4h up to UMAP creation. However, the total time including genome assembly is about 30h. Further research is thus needed to improve genome assembly speed.

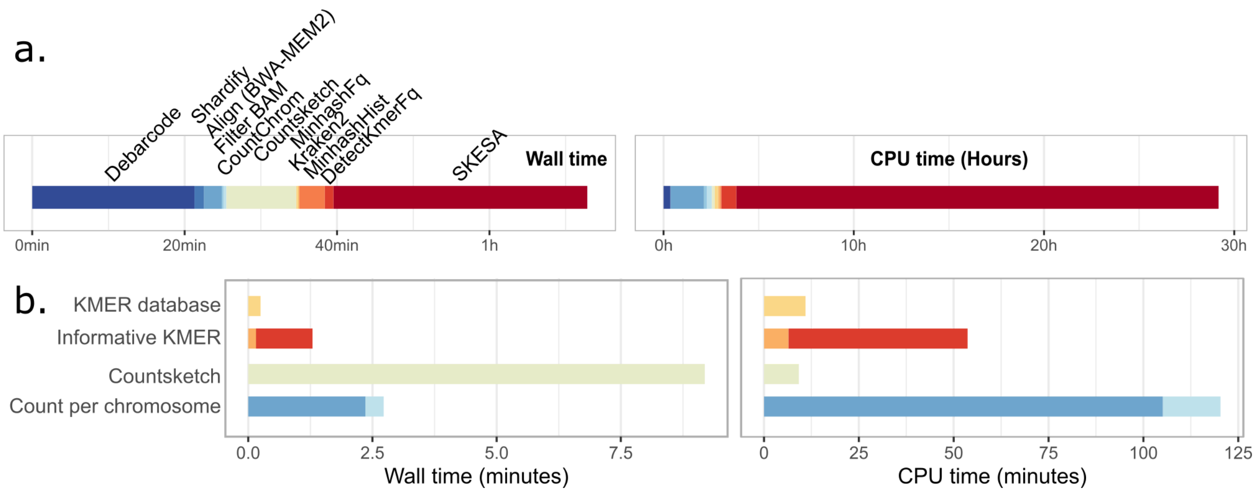

**Supplemental Note Figure 6: Wall time and CPU time for the entire Bascet/Zorn pipeline**, from raw reads to clustering analysis and assembled genomes. Benchmarked on the ATCC 10-strain mock community with PTA. **(a)** End-to-end time. **(b)** Time needed to produce different dimensional reductions.

### Bascet processing time versus CleanBar

CleanBar is the first published software specifically designed for cell barcode identification of Atrandi WGS-based libraries.[7] However, there are several key differences: **(1)** CleanBar is unable to read typical compressed FASTQ files for input and likewise does not write compressed output. **(2)** CleanBar writes one output file per barcode, which causes issues as described earlier in this document. **(3)** The CleanBar manuscript includes long-read data on libraries where library PCR was purposefully omitted, contrary to instructions from Atrandi. This results in a large number of reads with incomplete barcodes that cannot be assigned to a cell.

Properly debarcoding such reads is also more challenging, as it takes more time to process an incomplete read.

To enable benchmarking, we added support for the provided Atrandi WGS Pacbio raw FASTQ files to Bascet.[7] Thread settings for Bascet Debarcode were tuned, as we noted that compression of the output file is increasingly rate limiting in this setting (as there are fewer reads to debarcode, but more DNA sequence to compress).

We benchmarked against CleanBar (Github hash #a8fcaef) on their Pacbio dataset (NCBI SRA accession SRR31758484). Benchmarks include compression of the debarcoded reads for Bascet, while CleanBar does not offer compression. CleanBar also requires separate decompression of the input, which we have not included in the time. 40 threads were used for Bascet; CleanBar does not support multithreading. Benchmarking was performed on an Intel Xeon Gold 6138 CPU with 192GB RAM (30GB RAM given to Bascet).

In summary, Bascet is **9.2x** faster, despite performing additional work (compressed reading and writing; **Sup Note Fig 7**). We attribute the difference in speed, despite CleanBar being implemented in C, to several factors: **(1)** CleanBar does not support multithreading. **(2)** Bascet uses memory-mapped memory. **(3)** Bascet uses SIMD/SSE/AVX instructions in several places, making better use of modern hardware.

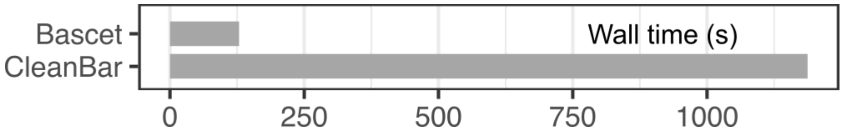

**Supplemental Note Figure 7: Comparison of Bascet/Zorn's speed to CleanBar.**

### Comparison to Mash

Mash is a common tool for genome comparison.[8] It uses min-hashes to first generate genome fingerprints; as the fingerprints are small, pairwise comparisons of the fingerprints are fast.

However, Mash has three problems that make it unsuitable for scMetaG analysis:

1. The software is unable to receive the reads of all cells in a few large files, and thus forces each genome to be stored in individual files. This is inappropriate as described in earlier sections.
2. Mash is based on MinHash as a distance measure between cells. MinHash approximates the Jaccard distance but has been shown to only be accurate for large sets of KMERS.[9] This can be compensated for using more hash functions, effectively keeping more of the data. However, there is still a lower limit for genome coverage when this breaks down.[9]
3. The tool is only capable of producing pairwise distances. This means that, for N genomes, N*(N-1) comparisons are required. In other words, the speed complexity is $O(N^2)$ (big-o notation[10]). While Mash reduces the time for pairwise comparison, the growth in time still renders this algorithm unsuitable for large numbers of genomes.

To get an idea of the time, we ran Mash v.2.3 on 10k simulated *Bacillus pacificus* genomes (results in **Sup Note Fig 8**; Intel Xeon Gold 6138 2.00GHz). Sketching was fast, with all genomes processed in 45 seconds (10 cores, 88 Mbp total sequence). However, computing pairwise distances ("mash triangle") required 13 min 53s. A scMetaG dataset of 1M cells would require $100^2$ more comparisons, in the order of 2300 hours (96 days). This duration does not include construction of the UMAP itself, which would best proceed by pruning the distance matrix to generate a kNN-graph. While a rewrite of Mash to use multiple cores and computers would significantly speed up the process, the $O(N^2)$ nature of the problem would still pose a challenge. The currently largest single-cell dataset has 100M cells,[11] and would require 2,600 years for Mash to process. With multithreading, this would still take over a year to complete, showing the infeasibility of the approach.

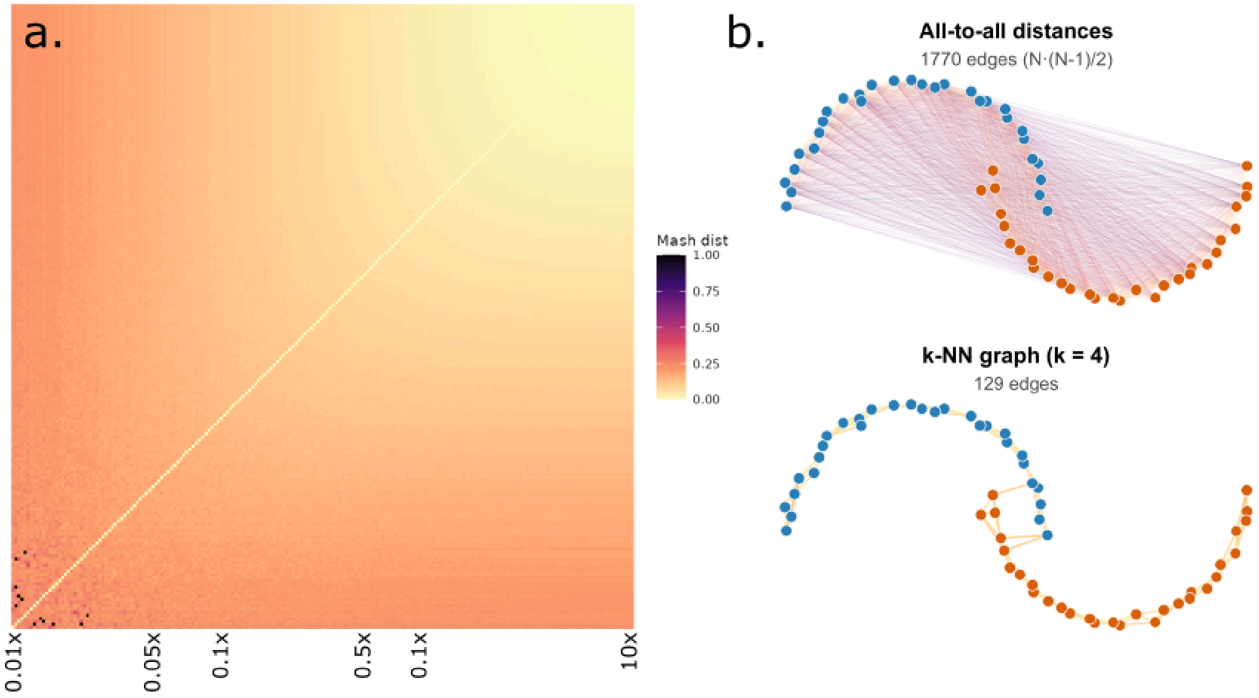

**Supplemental Note Figure 8: Comparison to Mash and MinHash distance.** (**a**) Mash distances between cells having the same genome, sampled uniformly at different coverage along a log scale. MinHash reports artificially high distance for low coverage genomes. (**b**) Mash stores all pairwise distances, while Bascet/Zorn computes the k-nearest neighbours, drastically reducing the amount of compute, while still finding the structure of the graph. CountSketch is better for approximating kNN graphs efficiently.

To bypass the need for pairwise comparisons, Zorn (together with Seurat) instead searches for the k-nearest neighbours using locality-sensitive hashing (LSH).[12] This effectively buckets similar cells in a manner that enables direct lookup of the k most similar cells. While the method is approximate, for sufficient k, this does not affect the overall topology of the kNN-graph.

Mash estimates genome size for use in its distance metric (https://mash.readthedocs.io/en/latest/distances.html). Because we expect this to be difficult for sparsely sampled genomes, Bascet/Zorn instead avoids the need for a genome size estimate. Instead, by using cosine distance,[13] the influence of genome size and sequencing depth is removed.

### Benchmarking of *k*-mer reduction methods

Single-cell analysis typically relies on the Leiden clustering method[14] to assign cell types.[15] We benchmarked its performance by comparing it to species-assignment by dominant taxonomy (here: which genome most reads align to). We used the Seurat implementation of Leiden clustering, which first requires a kNN-graph to be constructed. Two key parameters are k (number of neighbours in the approximate nearest neighbour graph) and resolution (a Leiden parameter, where higher resolution means more clusters). Grid search was performed on our simulated data and the ATCC 10-strain mock community PTA data (see **Online Methods**) to see which parameters were best (**Sup Note Fig 9**). While this is not possible on samples of unknown composition, it gives a qualitative idea of how to best set these parameters (in our experience, there is no correct parameter choice, and scRNAseq data is only interpreted qualitatively).

Because parameter choice affected both the number of clusters, and their purity (mixing of cells across species), we came up with reproducible but still qualitatively picked criteria for the best parameter choice: for a clustering with clusters ($C_j$) and alignment species labels ($S_i$), we computed cluster purity as $purity = \frac{1}{N}\sum_j max_i|c_j \cap S_i|$ , where $N$ is the total number of cells. We also recorded the largest-cluster fraction, $\frac{1}{N}max_j|c_j|$, and the minimum non-empty cluster size, $min|c_j|$. The target cluster number was the expected number of species, $K=10$. Candidate solutions were considered acceptable if purity was at least 0.7 and the largest-cluster fraction was at most 0.9. Among acceptable solutions with exactly $K$ clusters, we selected the clustering with highest purity, then lowest largest-cluster fraction, then largest minimum cluster size. If no exact-$K$ solution existed, we selected the solution with the least deviation from $K$.

From this analysis, we find that the alignment count table needs the least Leiden resolution, the Kraken2 and informative *k*-mer count table (100k KMERs) are roughly on par, and countsketch might need a bit higher resolution. This is expected, as aligned counts are the closest to the defined ground truth, Kraken2 has prior knowledge of which *k*-mers belong together, and countsketch keeps all *k*-mers at the cost of complex geometry in the reduced space. For the ATCC 10-strain mock community PTA data, not even the alignment workflow gives the correct number of clusters, which is in part to blame on some strains being hard to lyse (*D. radiodurans* and *C. beijerinckii*), and possibly also doublets/free-floating DNA.

In addition to clusterability by Leiden, another factor that may influence the choice of clustering method is the compute time (**Sup Note Fig 6b**). Kraken2 is the fastest option, while informative KMERs is the slowest on a single node. Countsketching can be the fastest option if multiple compute nodes are used. However, all of these methods are faster than counting reads aligning to different genomes.

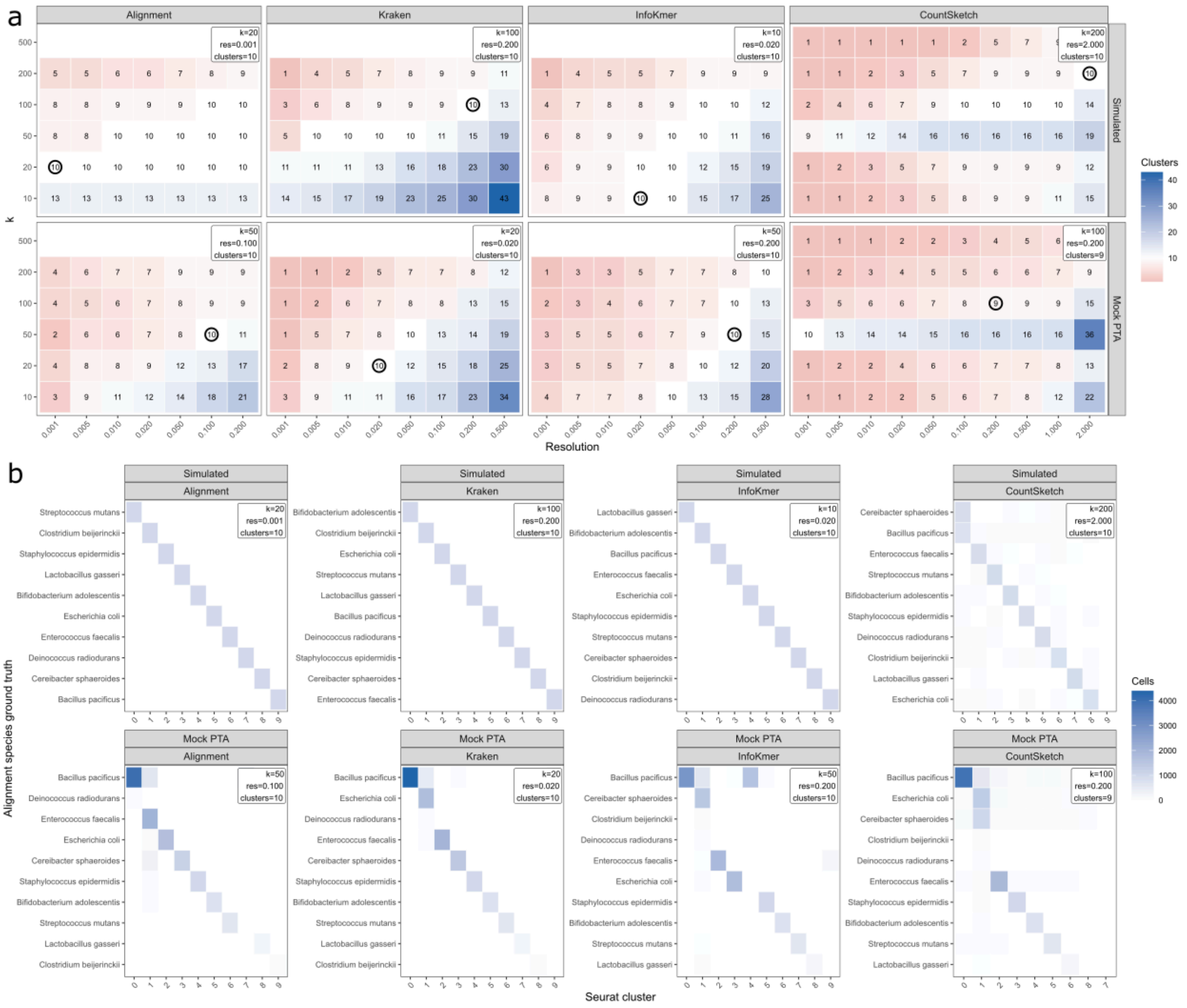

**Supplemental Note Figure 9: Benchmarking of *k*-mer reduction methods. (a)** Grid search for the best Leiden parameters. **(b)** Confusion plot of Leiden cluster *vs* species assigned by dominant taxonomy ID assignment.

### Analysis of Microbe-seq data from the human gut

Our Bascet/Zorn pipeline is designed to process any scMetaG data from any source. It can also take individually deposited cells or isolates and assemble them into TIRPs, ready for sharded processing. As an example, **Sup Note Fig 10** shows a UMAP of scMetaG data, generated from human fecal samples using the Microbe-seq method.[16] The processing time is entirely dominated by the download time of the reads of ~21,000 individual cells. Bascet/Zorn has support for parallel download of data from both SRA and NCBI.

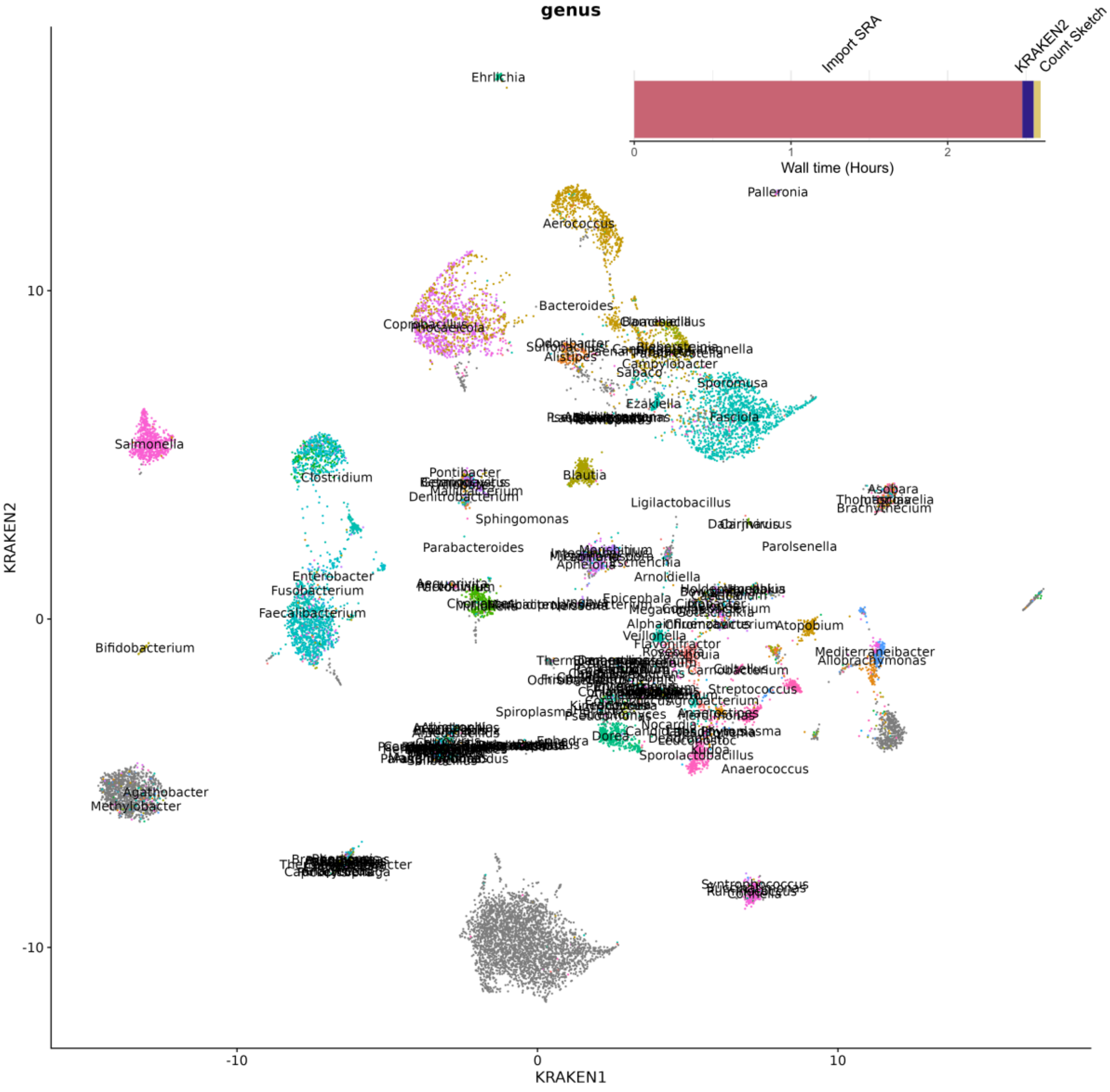

**Supplemental Note Figure 10:** UMAP of scMetaG data, generated from human fecal samples using Microbe-seq.[16]

### Doublet analysis can be applied to scMetaG data

To assess if typical scRNAseq doublet removal works, we used scDblFinder[17] and SingleCellExperiment[18] on the Kraken2 count matrix for the ATCC 10-strain mock community PTA data (**Sup Note Fig 11**). As expected, cells "between" clusters have high doublet scores. Thus, existing doublet detection methods can be used together with our Zorn R package.

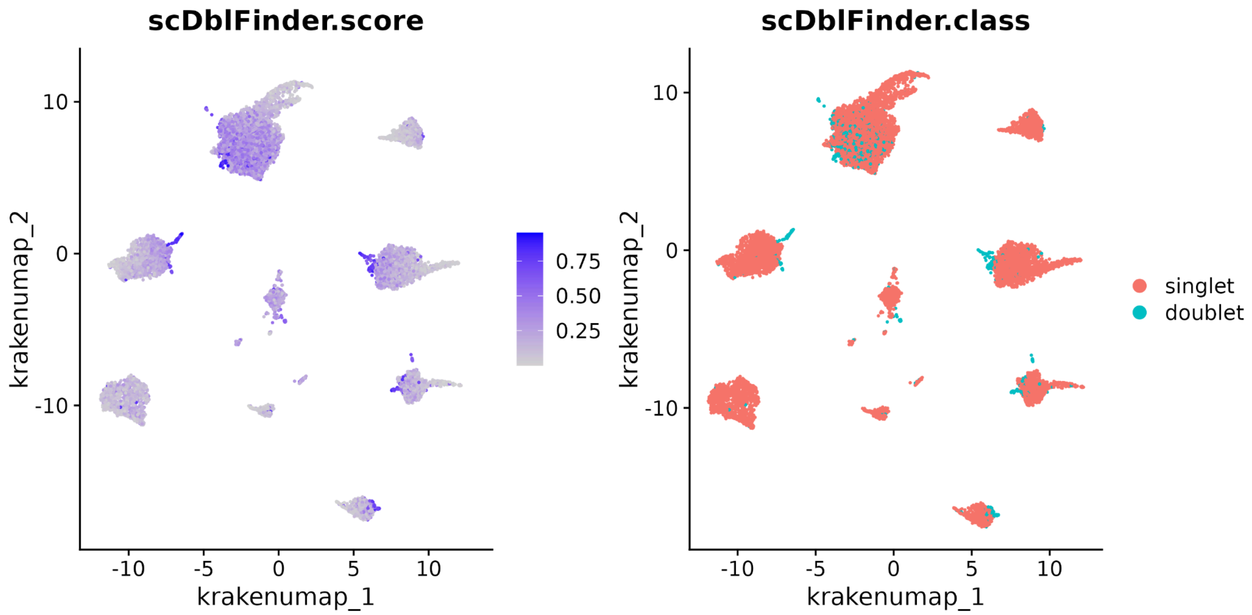

**Supplemental Note Figure 11:** scDblFinder[17] doublet analysis applied to the ATCC 10-strain mock community PTA data.

### Comparison of count normalization methods

There are common methods for preprocessing count table data before producing a UMAP, both available in Seurat and applicable to the output from Bascet (alignment count tables and informative *k*-mer count tables):

- RNA-seq analysis performs size factor normalization (divide by total counts), highly variable gene selection (comparison of variance to other genes at the same expression level), PCA to reduce dimensions, and UMAP.
- ATAC-seq analysis performs TF-IDF (Term Frequency-Inverse Document Frequency) normalization,[19] selection of top features (by how much they explain the data), SVD, and UMAP (commonly excluding the first SVD component).

We compared the two approaches on the ATCC 10-strain mock community PTA data (**Sup Note Fig 12**). Some clusters appear better with one approach but not the other. The online documentation for Bascet/Zorn gives code for both analyses as examples but with ATAC-seq as the default. Qualitatively, both approaches seem to function in practice, but we base our choice on two motivations: **(1)** Whole-genome sequencing is intuitively more similar to ATAC-seq (of DNA) than to analysis of RNA. **(2)** The TF-IDF transformation aims to weigh the importance of features based on their frequency. We reason that capture biases, such as by GC content, can cause certain features to be overrepresented and thus less informative. TF-IDF is a safe default choice to counteract this problem. We however typically do not see that the first SVD component reflects sequencing depth as it does for ATAC-seq (**Sup Note Fig 12**); this is to be expected, as depth is the primary component sought after in scMetaG analysis. Thus, the first SVD component should likely be kept. Note that this analysis is qualitative, and further research in the area of normalization is warranted.

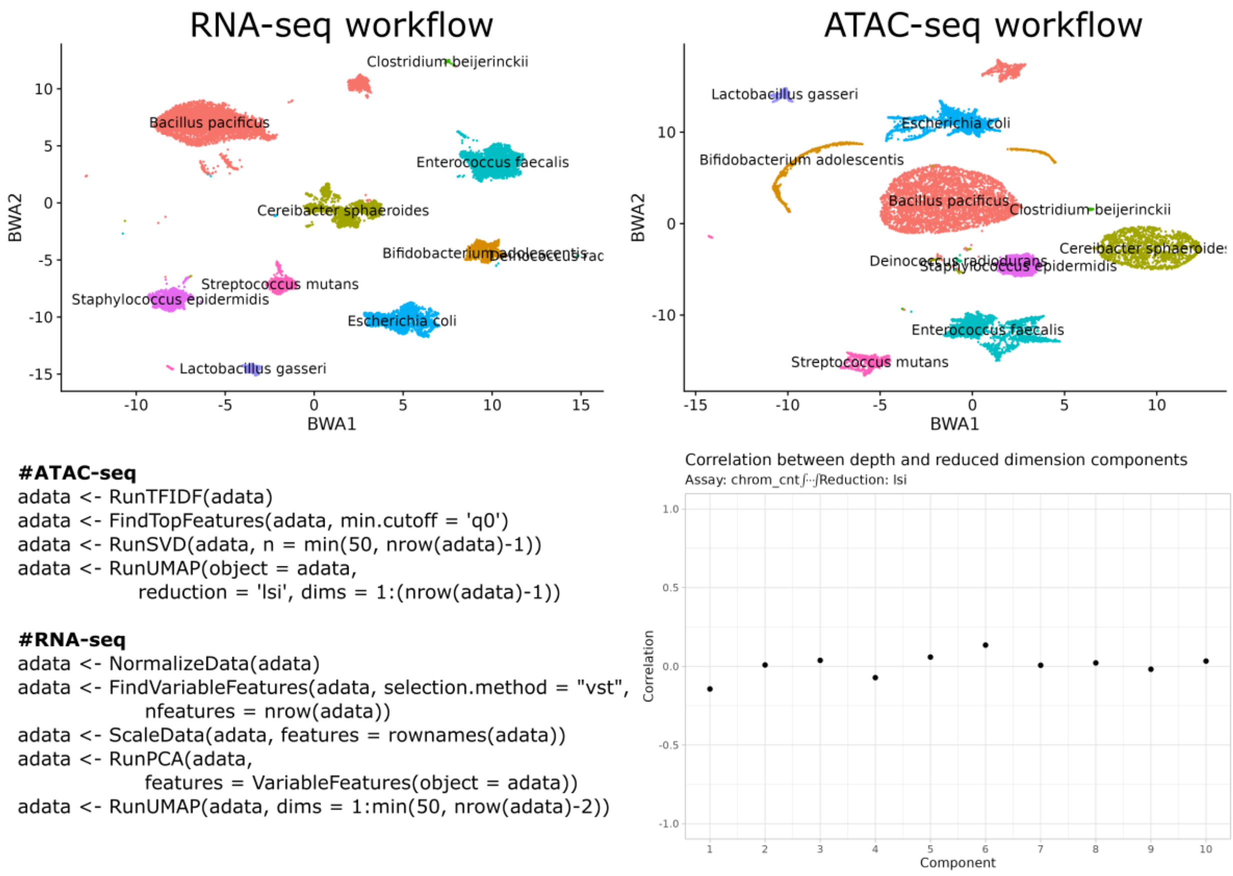

```
#ATAC-seq
adata <- RunTFIDF(adata)
adata <- FindTopFeatures(adata, min.cutoff = 'q0')
adata <- RunSVD(adata, n = min(50, nrow(adata)-1))
adata <- RunUMAP(object = adata,
          reduction = 'lsi', dims = 1:(nrow(adata)-1))

#RNA-seq
adata <- NormalizeData(adata)
adata <- FindVariableFeatures(adata, selection.method = "vst",
          nfeatures = nrow(adata))
adata <- ScaleData(adata, features = rownames(adata))
adata <- RunPCA(adata,
          features = VariableFeatures(object = adata))
adata <- RunUMAP(adata, dims = 1:min(50, nrow(adata)-2))
```

**Supplemental Note Figure 12:** Comparison of RNA-seq *vs* ATAC-seq based normalization. Example code for the normalizations is provided along with the Seurat DepthCorrelation QC.

### Power calculations

Single-cell analysis has for a long time been done for n=1 samples, with the aim of studying intrasample characteristics (e.g., finding new cell types) rather than comparing samples, or drawing conclusions from large cohorts. A few methods exist for power calculations, but they are not widely used in practice. Software for the purpose is listed in the following table (**Sup Note Table 1**).

| | Data distribution | Year of publication |
|---|---|---|
| powsimR[20] | Negative binomial | 2017 |
| scDesign[21] | A mixture of gamma and normal distributions on log-transform normalized data | 2019 |
| SCOPIT[22] | Poisson of Multinomal distribution | 2019 |
| POWSC[23] | A mixture of zero-inflated Poisson and log-normal Poisson distributions | 2020 |
| Hierarchicell[24] | Negative binomial | 2021 |
| scPower[25] | Negative binomial | 2021 |
| scDesign2[26] | Multifactor-based simulation | 2021 |
| scDesign3[27] | Multifactor-based simulation | 2024 |
| scPS[28] | Distribution-free | 2024 |

**Supplemental Note Table 1.** List of software for single-cell power calculation. Extended from a previous study.[28]

### Lysis protocol optimization and additional datasets

For initial lysis optimization, we first attempted 6 protocols (**M1-M6**), compared qualitatively using microscopy. Some protocols were further compared by qPCR using PCRBIO HS Taq Mix Red (Cat no. PB10.23-02; primer sequences in **Sup Note Table 2; Sup Note Fig 13**). The qPCR was performed with DNA inside SPCs, as well as in free solution. We then continued to optimize lysis using sequencing as a readout (**R1-R4; Sup Note Table 3** and **Sup Note Fig 14**). **R4** is our final optimized lysis protocol, described in the **Online Methods.**

**M1:** Based on a previous protocol[29] with custom modifications. Resuspend the SPCs in a PBS buffer with 50U/ul Lysozyme (Cat no. E-0057-D2, Biosearch Technologies), 22U/ml Lysostaphin (Cat no. L9043, Sigma), 250U/ml Mutanolysin (Cat no. SRE0007, Sigma) and incubate at 37°C overnight. The next day add 0.5mg/ml Achromopeptidase (Cat no. A3547, Sigma) followed by incubation at 37°C for 6-8 hours. Spin down and wash SPCs with 1X WB. Resuspend the SPCs in 1 mg/ml Proteinase K (P8107AA, NEB), 0.5% SDS, Incubate overnight at 40°C.

**M2:** Based on a previous protocol.[30] Resuspend the SPCs in a TE buffer containing: 2.5mM EDTA, 10mM NaCl, 5U Lysostaphin (Cat no. L9043, Sigma), 50U Mutanolysin (Cat no. SRE0007, Sigma), 20mg Lysozyme (Cat no. E-0057-D2, Biosearch Technologies) and incubate at 37°C overnight. The next day treat with TE buffer containing 4U Proteinase K (P8107AA, NEB), 1% Triton X 100 (Cat no. X100, Sigma), 100mM NaCl. Incubate at 55 °C for 30 minutes.

**M3:** Based on a previous protocol.[31] Resuspend the SPCs in Buffer 1: 20mM Tris HCl pH 8, 10mM EDTA, 100mM NaCl, 1% Triton X 100 (Cat no. X100, Sigma), 20mg/ml Lysozyme (Cat no. E-0057-D2, Biosearch Technologies) and incubate at 37°C for 1 hour. Treat with Buffer 2: Tris pH8, 20mM EDTA, 100mM NaCl, 1% SDS, 200ug/ml Proteinase K (P8107AA, NEB) and incubate at 55°C for 30 minutes.

**M4:** Based on a suggestion from Atrandi. Resuspend the SPCs in an Enzyme cocktail: 100U/ul Lysozyme (Cat no. E-0057-D2, Biosearch Technologies), 500U/ml Mutanolysin (Cat no. SRE0007, Sigma), 22U/ml Lysostaphin (Cat no. L9043, Sigma), 0.5mg/ml Achromopeptodase (Cat no. A3547, Sigma), and incubate at 37°C for 1 hour.

**M5:** Based on a suggestion from Atrandi. Resuspend the SPCs in the enzyme cocktail from M4 and incubate at 37°C for 1 hour. Spin down and wash SPCs with 1X WB. Resuspend the SPCs in 200ug/ml Proteinase K (Cat no. P8107AA, NEB), 1% SDS, 10mM EDTA, 10mM Tris-HCl pH7.5 and incubate at 55°C for 30 minutes. Next the SPCs were treated with the following lysis buffer: 0.8M KOH, 20mM EDTA, 200mM DTT (Cat no. P2325, Thermo Fisher Scientific) and incubated at RT for 15 minutes.

**M6:** Based on a previous protocol, modified from M2.[30] TE buffer containing: 2.5mM EDTA, 10mM NaCl, 5U Lysostaphin (Cat no. L9043, Sigma), 50U Mutanolysin (Cat no. SRE0007, Sigma), 20mg Lysozyme (Cat no. E-0057-D2, Biosearch Technologies) + 0.5mg/ml Achromopeptidase (Cat no. A3547, Sigma) at 37°C overnight. TE buffer with 4U Proteinase K (P8107AA, NEB), 1% Triton X 100 (Cat no. X100, Sigma), 100mM NaCl at 55 °C for 30 minutes.

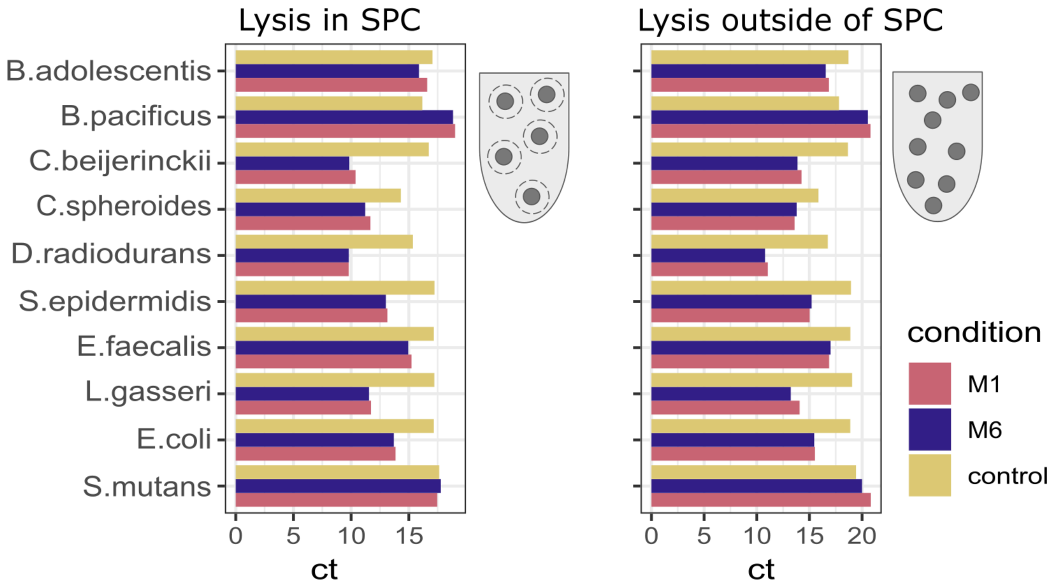

**Supplemental Note Figure 13:** *Verification of lysis for different species using qPCR.*

**Lysis optimization, validated by sequencing**

The SPCs were fixed in chilled 100% methanol before the initiation of the lysis. SPCs were divided into different tubes and treated with different lysis methods as below. Some, but not all protocols, were tested by sequencing. The results are shown **Sup Note Fig 14**.

**R1**: Adapted from Atrandi with custom modifications. SPCs were resuspended in buffer containing 100 U/μL Lysozyme, 0.2% Pluronic F-68, 200 mM NaCl, 2 mM EDTA, 20 mM Tris-HCl pH7.5 and incubated at 37°C for 30 min, 900 rpm shaking. SPCs were spun down and resuspended in 1mL of 200μg/mL Proteinase K, 1% SDS, 10 mM EDTA, 10 mM Tris-HCl pH7.5 and incubated at 50 °C for 30 minutes. SPSs were pelleted and resuspended in 500 μL of 1X Wash Buffer (WB (10 mM Tris- HCl (pH 7.5) with 0.1% Triton X-100 (Cat no. X100, Sigma) and mixed with 500 μL of 2X Atrandi lysis buffer consisting of 0.8 M KOH, 20 mM EDTA and 200 mM DTT (total reaction volume 1 mL) and incubated at RT for 15 minutes.

**R2:** Adapted from Atrandi with custom modifications. SPCs were resuspended in buffer containing 100 U/μL Lysozyme, 500U/ml Mutanolysin, 0.2% Pluronic F-68, 200 mM NaCl, 2 mM EDTA, 20 mM Tris-HCl pH7.5 and incubated at 37°C for 1 hour 800 rpm shaking. SPCs were pelleted and resuspended in 1mL of 200μg/mL Proteinase K, 1% SDS, 10 mM EDTA, 10 mM Tris-HCl pH7.5 and incubated at 55°C for 30 minutes. SPCs were pelleted and resuspended in 500μL of 1X WB and mixed with 500 μL of 2X Atrandi lysis buffer and incubated at RT for 15 minutes.

**R3:** SPCs were resuspended in TE buffer containing: 2.5mM EDTA, 10mM NaCl, 5U Lysostaphin (Cat no. L9043, Sigma), 50U Mutanolysin (Cat no. SRE0007, Sigma), 20mg Lysozyme (Cat no. E-0057-D2, Biosearch Technologies) and 0.5mg/ml Achromopeptidase (Cat no. A3547, Sigma) and incubated at 37°C overnight at 800 rpm shaking. SPCs were pelleted and resuspended, washed in 1X WB 3 times and resuspended in 1 mL of 4U Proteinase K, 1%Triton X 100 and 100mM NaCl and incubated at 55°C for 30 minutes–1h at 800 rpm shaking.

**R4:** This protocol was used for the datasets presented in the main text. See **Online methods**

| | Target bacteria | Primer sequence | Target gene | Amplicon size (bp) | Reference |
|---|---|---|---|---|---|
| 1 | *Clostridium beijnerinckii* | CbeF- TGACACGATTTTTCATTCTCCA<br>CbeR- TCCATTGCCTTAATGACAGGT | *nifH* | 448 | 32 |
| 2 | *E. coli* | EcoF- CGTGGTGATTGATGAAACTG<br>EcoR- TGATACATATCCAGCCATGC | *uidA* | 564 | 33 |
| 3 | *Lactobacillus gasseri* | LgasF- ATCACATTCAACTCTCGCTG<br>LgasR- TCATTCATCTTCATCGTCCT | Locus tag: LGAS_0517 | 400 | 34 |
| 4 | *Staphylococcus epidermidis* | SepiF- GATATTCGCGATGAACTTGC<br>SepiR- ATCAGGTGTTGCAAATAGGG | Fibrinogen-binding protein | 414 | 35 |
| 5 | *Streptococcus mutans* | SmutF- TCGCGAAAAAGATAAACAAACA<br>SmutR- GCCCCTTCACAGTTGGTTAG | Fibrinogen-binding protein<br>Species specific locus | 479 | 36 |
| 6 | *Rhodobacter sp./Ceribacter sp.* | Cer_2-F: ATGTTGGACCTCCGCAAAG<br>Cer_2-R: TGCCATCTGATGCCGTATTG | (locus_tag="Rsph17029_0004"-Hypothetical protein) | 341 | |
| 7 | *Enterococcus faecalis* | Efa_2-F: AGTACCATTCGTGCCAGTTT<br>Efa_2-R: GCGTATTCTTGCGCTTGATG | *ddl*<br>D-Alanine-D-Alanine Ligase) | 399 | 37 |
| 8 | *Deinococcus radiodurans* | Dra_2-F: GAAGTCGAGGTGGCGTTTAT<br>Dra_2-R: TCTTGCCGATCTTGGGATTT | *gyrB* | 361 | 38 |
| 9 | *Bifidobacterium adolescentis* | BadF: GGTGATTACGCAGCATCCTT<br>BadR: CTTCCTCACAAACGTCAGCA | | | 39 |
| 10 | *Bacillus pacificus (cereus)* | BcerF: GTGGTTCTGCTGTATCTA<br>BcerR: CAGCACCAGTAACGTTTA | *entFM* | 183 | 40 |

**Supplemental Note Table 2.** List of primers used for qPCR.

| Protocol | Dataset |
|---|---|
| R1 | mock1_MDA |
| R2 | mock2_MDA |
| R3 | mock3_MDA<br>saliva1_MDA |
| R4 | mock4_MDA<br>mock5_PTA<br>saliva2_MDA |

**Supplemental Note Table 3.** List of sequenced datasets and corresponding protocols.

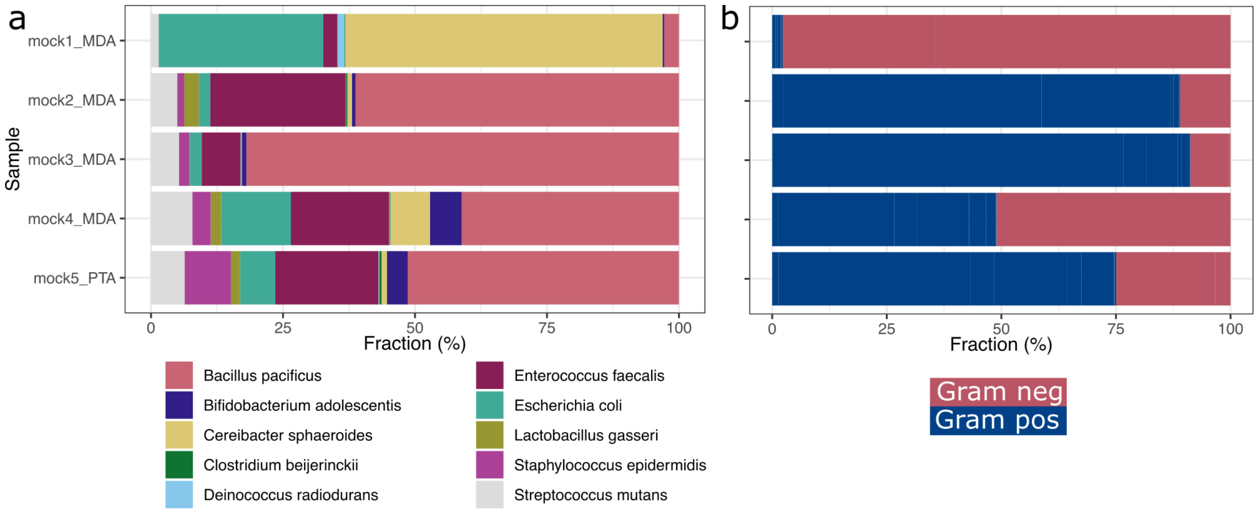

**Supplemental Note Figure 14.** Comparison of lysis protocols across all sequenced samples. **(a)** Fraction of reads per species. **(b)** Gram positive/negative fraction, normalized by the number of Gram positive/negative strains present in the pool. Mock samples 1-3 were treated using commercial protocols, while mock samples 4-5 used our custom protocols, developed in this study.

### Upstream software versions

Bascet integrates several software packages via Rust translation, both to simplify distribution and to speed up the code (described previously[41]). These are the versions of the software used for the results in this manuscript:

- BWA-MEM2, crates.io, bwa-mem2-pure-rs 0.2
- GECCO, crates.io, gecco, 0.5.4
- FASTQC, crates.io, fastqc-compliant-rs 0.4.2
- Kraken2, crates.io, kraken2-pure-rs 0.2.0
- Minimap2, crates.io, minimap2-pure-rs 0.5
- SKESA, crates.io, skesa-rs 0.2
- STAR, https://github.com/henriksson-lab/star-rs

For other upstream versions, see Cargo.toml in our repository (Bascet #c378038, v0.0.2).

### Cost comparison

##### Price breakdown for SPC-based single-cell metagenomics

| Kit/Reagent | Cost (SEK) | Reactions | Cost/rxn (SEK) |
|---|---|---|---|
| **From Atrandi** | | | |
| C2-chip (single) | 2300 | 6 | 383 |
| Barcoding Kit | 10900 | 3 | 3633 |
| Encapsulation kit | 6690 | 3 | 2230 |
| **Own Lysis** | | | |
| Ready-Lyse Lysozyme Solution 4 000 000 U | 4770 | 500 | 10 |
| Mutanolysin | 5194 | 500 | 10 |
| Achromopeptidase | 2121 | 500 | 4 |
| Lysostaphin | 6854 | 500 | 14 |
| Proteinase K | 400 | 100 | 4 |
| **Own MDA** | | | |
| Equiphi29 250U | 673 | ~10-20 | **260** |
| T4 polymerase | 1040 | ~20 | |
| T7 endonuclease | 1150 | ~20 | |
| DNA Polymerase Large Fragment (Klenow) - 200 units | 1030 | ~20 | |
| **Adapted PTA** | | | |
| ResolveDNA Whole Genome Amplification Kit v2.0 | 11042 | 20 | 552 |

| **Sequencing using novaseq 1.5B (one lane)** | |
|---|---|
| 19000 | for 10k cells |
| 1.9 | SEK/cell (10k) |

| **Library prep cost PTA** | |
|---|---|
| 11042 | total cost SEK |
| 1.1 | SEK/cell (10k) |
| 3.0 | SEK/cell with seq (10k) |
| 0.270 | eur/cell |

| **Library prep cost MDA** | |
|---|---|
| 6548 | total cost SEK |
| 0.7 | sek/cell (10k) |
| 2.6 | sek/cell with seq (10k) |
| 0.230 | eur/cell |

**Price breakdown multiwell-plate-based single cell metagenomics**

| **Takara picoplex** | |
|---|---|
| 12642 | eur for 480rxn |
| 26 | sek/cell |

| **Library prep using illumina DNA prep** | |
|---|---|
| Illumina DNA Prep, (M) Tagmentation (96 Samples, IPB) | |
| 31460 | sek for 96 |
| 327.7 | sek/cell |
| Illumina DNA/RNA UD Indexes Set A, Tagmentation (96 Indexes, 96 Samples) | |
| 4729 | sek for 96 |
| 49.3 | sek/cell |
| **Sequencing using novaseq 1.5B (one lane)** | |
| 19000 | for 384 cells |
| 49.5 | sek/cell |

| | |
|---|---|
| 403.0 | sek/cell for library prep (10k) |
| 49.5 | sek/cell for sequencing (10k) |
| 452.4 | sek/cell total |
| 40.720 | eur/cell |

**Cost ratio PTA: 151x**

Lower costs for SPCs can be achieved by using larger Novaseq flow cells (i.e. 25B).
